# Spatial close-kin mark-recapture methods to estimate dispersal parameters and barrier strength for mosquitoes

**DOI:** 10.1101/2025.05.11.653364

**Authors:** John M. Marshall, Shuyi Yang, Jared B. Bennett, Igor Filipović, Gordana Rašić

**Affiliations:** Divisions of Biostatistics and Epidemiology, School of Public Health, University of California, Berkeley, CA, USA; Innovative Genomics Institute, University of California, Berkeley, CA, USA; Mosquito Genomics, QIMR Berghofer Medical Research Institute, Brisbane, Australia

## Abstract

Close-kin mark-recapture (CKMR) methods have recently been used to infer demographic parameters for several aquatic and terrestrial species. For mosquitoes, the spatial distribution of close-kin pairs has been used to estimate mean dispersal distance, of relevance to vector-borne disease transmission and genetic biocontrol strategies. Close-kin methods have advantages over traditional mark-release-recapture (MRR) methods as the mark is genetic, removing the need for physical marking and recapturing that may interfere with movement behavior. Here, we extend CKMR methods to accommodate spatial structure alongside life history for mosquitoes and comparable insects. We derive kinship probabilities for parent-offspring and full-sibling pairs in a spatial context, where an individual in each pair may be a larva or adult. Using the dengue vector *Aedes aegypti* as a case study, we use an individual-based model of mosquito life history to test the effectiveness of this approach at estimating parameters such as mean dispersal distance, daily staying probability, and the strength of a barrier to movement. Considering a simulated population of 9,025 adult mosquitoes arranged on a 19-by-19 grid, we find the CKMR approach provides unbiased and precise estimates of mean dispersal distance given a total of 2,500 adult females sampled over a three-month period using 25 traps evenly spread throughout the landscape. The CKMR approach is also able to estimate parameters of more complex dispersal kernels, such as the daily staying probability of a zero-inflated exponential kernel, or the strength of a barrier to movement, provided the magnitude of these parameters is greater than 0.5. These results suggest that CKMR provides an insightful characterization of mosquito dispersal that is complementary to conventional MRR methods.

**Author summary:** Close-kin mark-recapture (CKMR) is a genetic analogue of mark-release-recapture (MRR) in which the frequency of marked individuals in a sample is used to infer demographic parameters such as census population size and mean dispersal distance. These methods have been widely applied to aquatic species; however their application to mosquitoes is yet to be rigorously explored. Previous theoretical work demonstrated the potential for CKMR to infer parameters such as population size and mortality rate for randomly-mixing mosquito populations, and close-kin-based methods have been used to infer movement patterns for *Aedes aegypti* mosquitoes in Singapore and Malaysia. Here, we use simulations to explore the potential for formal CKMR methods to characterize mosquito dispersal patterns. We find that formal CKMR methods are able to accurately estimate mean dispersal distance, and to estimate additional parameters, such as the strength of a landscape barrier and the probability that a mosquito remains within its population node each day. CKMR and other close-kin-based methods provide insights into mosquito dispersal complementary to commonly-used alternatives such as MRR, as they capture displacement across several generations and are not compromised by the marking process.

## 1. Introduction

Malaria, dengue, chikungunya and other mosquito-borne diseases continue to pose a major burden throughout much of the world [1, 2]. Novel biological and genetics-based interventions, such as releases of mosquitoes infected with *Wolbachia* or engineered with gene drives, offer much promise to complement traditional control tools such as insecticide-treated nets, vaccines and antimalarial drugs. A common feature of these novel tools is the need for a detailed understanding of mosquito movement in order to design effective field trials and interventions, and to address biosafety concerns. In a recent randomized controlled trial (RCT) of *Wolbachia*-based population replacement of the dengue vector, *Aedes aegypti*, in Yogyakarta, Indonesia, *Wolbachia* was observed to spread significantly from intervention to control areas within one year of release [3]. This highlights the importance of quantifying mosquito movement to determine optimal spatial units for vector control RCTs. Predicting intentional geographic spread of self-propagating interventions such as *Wolbachia* and gene drive is also crucial, as is assessing the potential for confinement and logistics of reversibility during a trial [4].

A handful of methods are available to characterize mosquito movement patterns. The most direct of these is mark-release-recapture (MRR), hundreds of which studies have been conducted for *Ae. aegypti* and the malaria vector *Anopheles gambiae* in recent decades [5]. In MRR, a portion of a population is captured, marked and released, and subsequent collections are checked for recaptures. The fraction of recaptures over time can be used to infer population size and daily mortality, while times and distances between release and recapture events can be used to infer dispersal patterns. A major shortcoming of MRR is that inferred dispersal patterns may be modified by the process of marking and capturing, and for mosquitoes specifically, releasing females may increase the risk of local disease transmission. Several genetic methods are available to characterize mosquito movement on a larger spatial scale, effectively estimating dispersal averaged over several generations. Wright’s fixation index, *F*_*ST*_, can be calculated using genetic markers such as single nucleotide polymorphisms or microsatellites [6, 7], and population assignment tests can be used to infer movement between well-structured populations at a scale beyond the mean dispersal range of a species [8].

Here, we theoretically explore the role that close-kin mark-recapture (CKMR) methods could play in characterizing the dispersal patterns of *Ae. aegypti* mosquito vectors. In CKMR, the detection of a close-kin pair (parent-offspring, siblings, etc.) in a sample is analogous to the recapture of a marked individual in the MRR method [9]. Detection of several close-kin pairs separated by a given distance provides information about movement that occurred over a small number of generations, thus informing dispersal patterns on an intermediate spatial scale, between that of MRR methods and most other genetic approaches. Advantages of CKMR methods stem from the mark being a genetically-inferred close-kin relationship, removing the need for physical marking and recapturing. Two recent studies have used the spatial distribution of close-kin pairs to characterize *Ae. aegypti* movement patterns [10, 11], both using approaches inspired by the CKMR formalism [9]. Jasper *et al*. [10] estimate relatedness across three orders of kinship and estimate life stage-specific dispersal by considering possible life history events between each kinship pair. Filipović *et al*. [11] consider up to three orders of kinship and use coordinates of close-kin pairs to infer the distribution of distances between the birth and ovipositing sites of breeding females, and fit these distances to a variety of dispersal kernels.

Here, we apply formal CKMR methods with spatial structure to estimate movement parameters of *Ae. aegypti*. Formal CKMR methods are based on explicit calculations of kinship probabilities, and the likelihood of observing a given number and category of close-kin pairs across space and time [9]. While the availability of spatial kinship information has been discussed as having the potential to inform dispersal parameters within a formal CKMR framework [9, 12, 13], very few studies have explored this through simulation [14, 15]. Our approach builds upon CKMR methods specific to the life history of mosquitoes, previously used to estimate demographic parameters such as census population size, and adult and larval mortality rates [16]. As in that study, we use an *in silico* model of mosquito life history, this time with spatial structure, to generate kinship data and validate our inference methods. Using this approach, we determine optimal sampling schemes (sampled life stages, sample size and spatial distribution of traps) to accurately and efficiently estimate dispersal parameters, including mean dispersal distance, daily staying probability, and the strength of a barrier to movement.

## 2. Methods

### 2.1. Mosquito life history and spatial structure

As per our previous mosquito CKMR study [16], we base our analysis on a discrete-time version of the lumped age-class model [17, 18], applied to mosquitoes [19] (Figure 1A). This model considers discrete life history stages - egg (E), larva (L), pupa (P) and adult (A) - with sub-adult stages having defined durations - *T*_*E*_, *T*_*L*_ and *T*_*P*_ for eggs, larvae and pupae, respectively. We use a daily time-step, since mosquito samples tend to be recorded by day, and this is adequate to model the organism’s population dynamics [20]. Daily mortality rates vary according to life stage - *µ*_*E*_, *µ*_*L*_, *µ*_*P*_ and *µ*_*A*_ for eggs, larvae, pupae and adults, respectively - and density-dependent mortality occurs at the larval stage. Sex is modeled at the adult stage - half of pupae emerge as females (F), and the other half as males (M). Females mate once upon emergence, and retain the genetic material from that mating event for the remainder of their lives. Males mate at a rate equal to the female emergence rate which, for a population at equilibrium, is equal to the female mortality rate, *µ*_*A*_. Females lay eggs at a rate, *β*, which is assumed to be independent of age.

**Fig 1.**
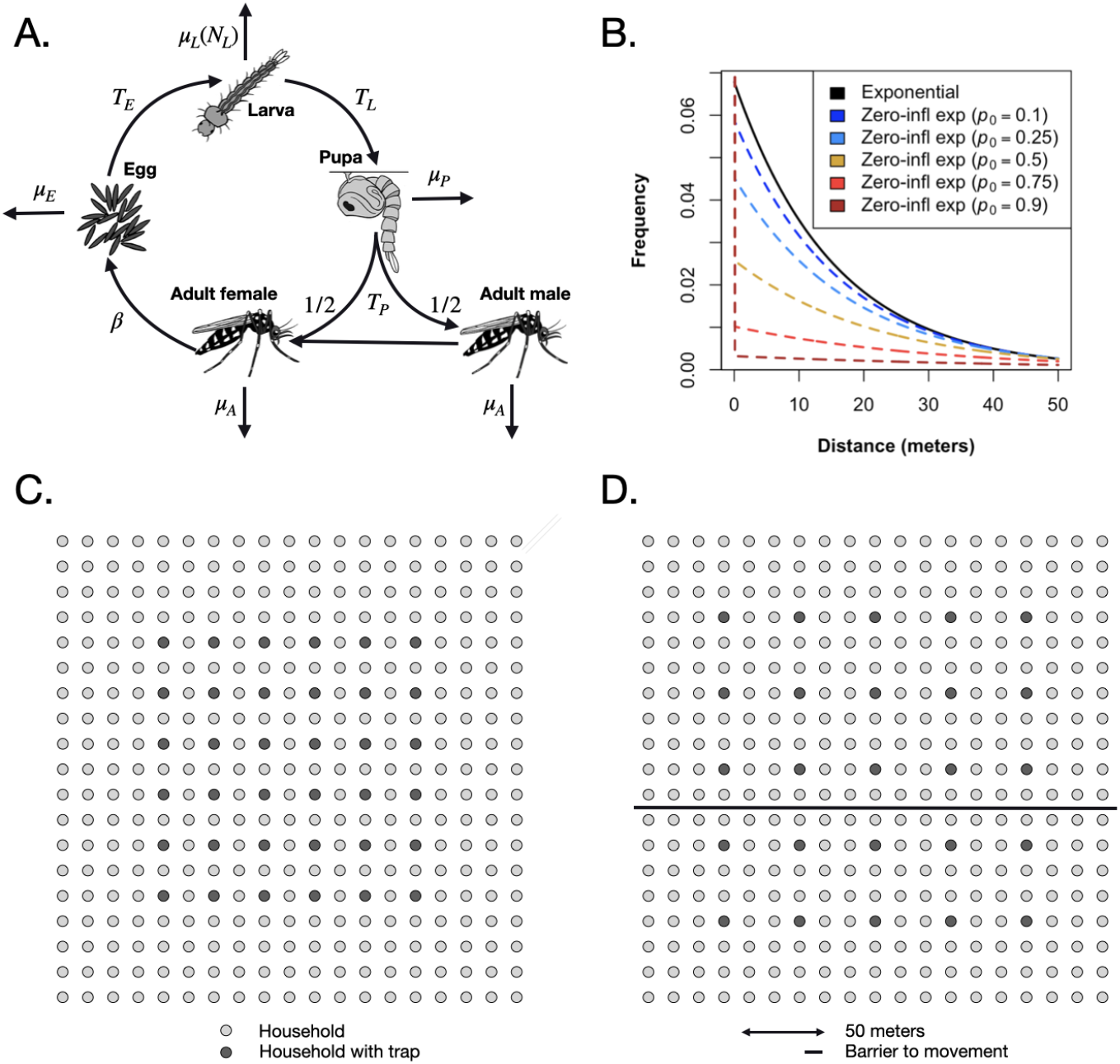
Mosquito life history and spatial structure. In the lumped age-class model, mosquitoes are divided into four life stages: egg, larva, pupa and adult **(A)**. The durations of the sub-adult stages are *T*_*E*_, *T*_*L*_ and *T*_*P*_ for eggs, larvae and pupae, respectively. Sex is modeled at the adult stage, with half of pupae developing into females and half developing into males. Daily mortality rates vary by life stage - *µ*_*E*_, *µ*_*L*_, *µ*_*P*_ and *µ*_*A*_ for eggs, larvae, pupae and adults, respectively. Density-dependent mortality occurs at the larval stage and is a function of the total number of larvae, *N*_*L*_. Females mate once upon emergence, and retain the genetic material from that mating event for the remainder of their lives. Males mate at a rate equal to the female emergence rate. Females lay eggs at a rate, *β*. In the spatial extension of the lumped age-class model, mosquito populations are distributed in space, with movement between them defined by an exponential (solid line) or zero-inflated exponential dispersal kernel (dashed lines) **(B)**. The daily probability of remaining in the same population, *p*_0_, is varied while preserving the mean dispersal distance. This value is trimmed from the plot, but specified in the key. Mosquito populations are distributed according to a 19-by-19 grid of households (circles), with mosquito traps distributed in select households (black circles) according to the sampling scheme **(C)**. In some simulations and analyses, a barrier to movement is included (solid line) **(D)**.

In extending the lumped age-class model to space, we consider a spatial distribution of mosquito populations, with movement between them defined by a dispersal kernel (Figure 1B-D). Discrete populations in the resulting metapopulation are considered to be randomly mixing populations to which the lumped age-class model applies. The resolution of the individual populations (in terms of size) should be chosen according to the dispersal properties of the species being considered. For *Ae. aegypti*, populations on the scale of households may be appropriate, as this species tends to disperse locally, often remaining in the same household for the duration of their lifespan [21]. For *An. gambiae*, dispersal occurs over larger distances and villages may be an appropriate population unit [6]. By default in this paper, daily per-capita movement probabilities between populations are derived from an exponential dispersal kernel (Figure 1B). For populations *i* and *j* a distance *d*_*ij*_ apart, the rate of movement between them is:

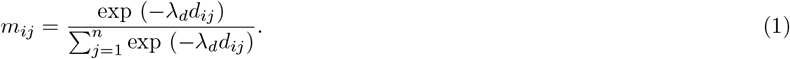

Here, 1/*λ*_*d*_ represents the mean daily dispersal distance, conditional upon movement, and *n* represents the number of populations in the landscape. For a given origin, *i*, the dispersal kernel entries, *m*(*i*,), sum to 1. Computing *m*_*ij*_ for all combinations of origins and destinations produces the movement matrix, *M*.

We also consider a zero-inflated exponential kernel (Figure 1B) which includes an additional parameter, *p*_0_, to represent the daily probability that a mosquito remains in the same population. For this kernel, we have:

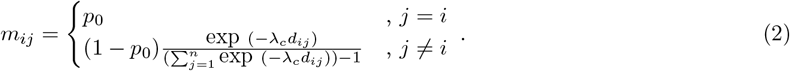

Here, 1*/λ*_*c*_ represents the mean daily dispersal distance, conditional upon movement, and the diagonal elements of the movement matrix, *m*_*ii*_, equal *p*_0_.

Default life history, demographic and dispersal parameters for *Ae. aegypti* are listed in Table 1. Given the difficulty of measuring juvenile stage mortality rates in the wild, these are chosen for consistency with observed population growth rates in the absence of density-dependence (see Sharma *et al*. [16] for formulae and derivations). Larval mortality increases with larval density and, according to the lumped age-class model, reaches a set value when the population is at equilibrium. Although mosquito populations vary seasonally, we assume a constant adult population size, *N*_*A*_, for this CKMR analysis, and restrict sampling to a period of three months, corresponding to a season. Minor population size fluctuations occur in the simulation model due to sampling and stochasticity. For dispersal, we consider movement within a 19-by-19 grid of households, based on a suburban setting such as Cairns in Queensland, Australia [21]. Mosquito traps are distributed in select households according to a specified sampling scheme (Figure 1C). In some simulations and analyses, a barrier to movement is included, which could represent a road, freeway or open park space (Figure 1D).

**Table 1.**
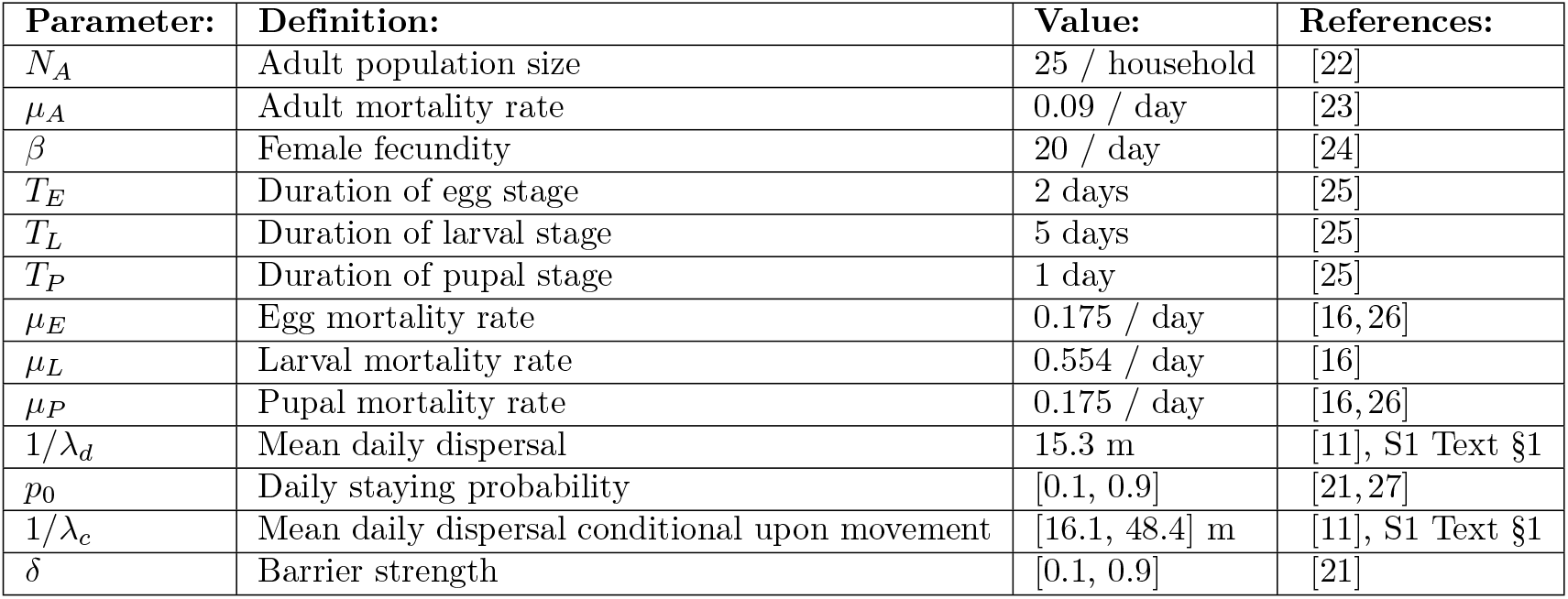
Demographic, life history and dispersal parameters for *Aedes aegypti* mosquitoes.

| Parameter: | Definition: | Value: | References: |
| --- | --- | --- | --- |
| $N_A$ | Adult population size | 25 / household | [22] |
| $\mu_A$ | Adult mortality rate | 0.09 / day | [23] |
| $\beta$ | Female fecundity | 20 / day | [24] |
| $T_E$ | Duration of egg stage | 2 days | [25] |
| $T_L$ | Duration of larval stage | 5 days | [25] |
| $T_P$ | Duration of pupal stage | 1 day | [25] |
| $\mu_E$ | Egg mortality rate | 0.175 / day | [16, 26] |
| $\mu_L$ | Larval mortality rate | 0.554 / day | [16] |
| $\mu_P$ | Pupal mortality rate | 0.175 / day | [16, 26] |
| $1/\lambda_d$ | Mean daily dispersal | 15.3 m | [11], S1 Text §1 |
| $p_0$ | Daily staying probability | [0.1, 0.9] | [21, 27] |
| $1/\lambda_c$ | Mean daily dispersal conditional upon movement | [16.1, 48.4] m | [11], S1 Text §1 |
| $\delta$ | Barrier strength | [0.1, 0.9] | [21] |

### 2.2. Kinship probabilities

Following the CKMR methodology of Bravington *et al*. [9] and its application to mosquito populations by Sharma *et al*. [16], we now derive spatial kinship probabilities for mother-offspring and full-sibling pairs based on the lumped age-class mosquito life history model. Each kinship probability is calculated as the reproductive output consistent with that relationship divided by the total reproductive output of all adult females in that population. In each case, we consider two individuals (adult or larva) sampled at known locations, *x*_1_ and *x*_2_, and times, *t*_1_ and *t*_2_, with probability symbols and references to equations listed in Table 2. Note that mosquito sampling is lethal.

**Table 2.**
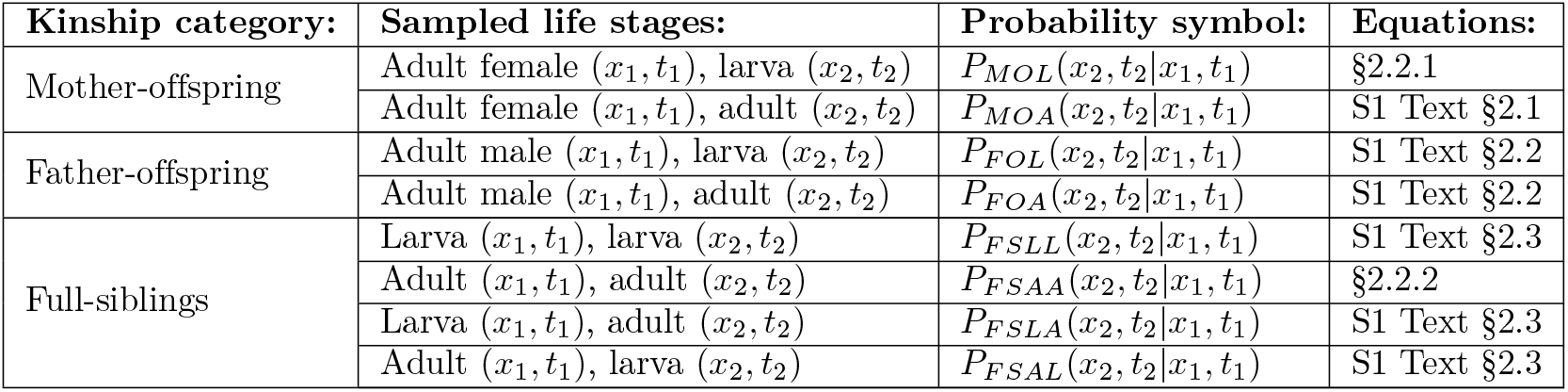
Kinship categories, sampled life stages, sampling times, locations, and probability symbols used in spatial close-kin mark-recapture analysis.

#### 2.2.1. Mother-offspring

Let us begin with the simplest possible kinship probability, *P*_*MOL*_(*x*_2_, *t*_2_|*x*_1_, *t*_1_), which represents the probability that, given an adult female sampled at location *x*_1_ on day *t*_1_, a larva sampled at location *x*_2_ on day *t*_2_ is her offspring. This can be expressed as the relative larval reproductive output at location *x*_2_ on day *t*_2_ of an adult female sampled at location *x*_1_ on day *t*_1_:

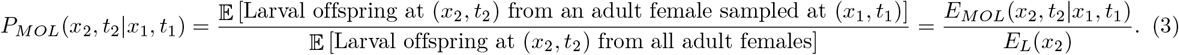

Here, *E*_*MOL*_(*x*_2_, *t*_2_|*x*_1_, *t*_1_) represents the expected number of surviving larval offspring at location *x*_2_ on day *t*_2_ from an adult female sampled at location *x*_1_ on day *t*_1_, and *E*_*L*_(*x*_2_) represents the expected number of surviving larval offspring at location *x*_2_ from all adult females in the population at times consistent with the time of larval sampling. Note that, since we are assuming a constant population size, *E*_*L*_(*x*_2_) is independent of time and is given by:

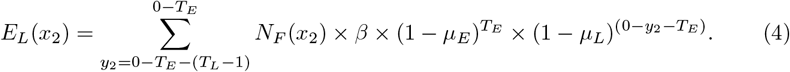

Here, *N*_*F*_ (*x*_2_) represents the equilibrium adult female population size, and *y*_2_ represents the day of egg-laying. Considering day 0 as the reference day (in place of *t*_2_), the egg must have been laid between days (0 − *T*_*E*_ − (*T*_*L*_ − 1)) and (0 − *T*_*E*_). Equation 4 therefore represents the expected number of offspring laid by all adult females at location *x*_2_ that survive the egg and larva stages up to the time of sampling (day 0).

*E*_*MOL*_(*x*_2_, *t*_2_|*x*_1_, *t*_1_), on the other hand, is specific to the sampled adult female, the location, *x*_2_, and the day of larval sampling, *t*_2_. This is given by:

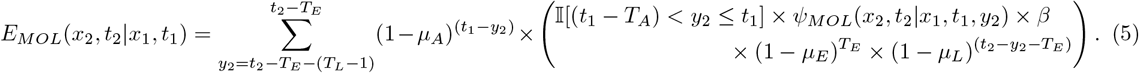

Here, the day of egg-laying, *y*_2_, is summed over days (*t*_2_ − *T*_*E*_ − (*T*_*L*_ − 1)) through (*t*_2_ −*T*_*E*_), for consistency with the larva being present on the day of sampling (Figure 2). The first term in the summation represents the probability that the adult female sampled on day *t*_1_ is alive on the day of egg-laying (*y*_2_), and the second term (in large brackets) represents the expected surviving larval output of this adult female at location *x*_2_ on day *t*_2_. This latter term is equal to the probability that the larval offspring is sampled at location *x*_2_ on day *t*_2_ given the mother is sampled at location *x*_1_ on day *t*_1_ and the egg is laid on day *y*_2_, *ψ*_*MOL*_(*x*_2_, *t*_2_ | *x*_1_, *t*_1_, *y*_2_), multiplied by their daily egg production, *β*, multiplied by the proportion of eggs that survive the egg and larva stages from the day they were laid up to the day of sampling. An indicator function is included to limit consideration to cases where the day of egg-laying lies within the adult female’s possible lifetime - i.e., between days *t*_1_ and (*t*_1_ − *T*_*A*_), where *T*_*A*_ represents the maximum possible age of an adult mosquito. Although adult lifetime is exponentially-distributed, a value of *T*_*A*_ may be chosen that captures most of this distribution and leads to accurate parameter inference.

**Fig 2.**
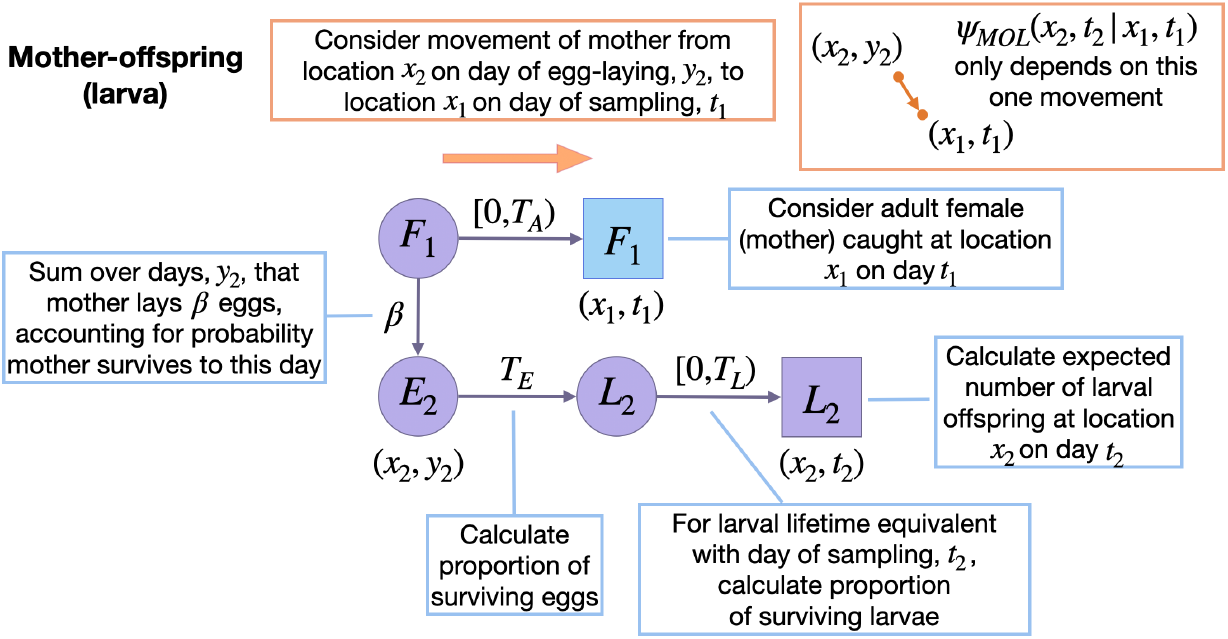
Schematic representation of spatial mother-larval offspring kinship probability. Parameters and state variables are as defined in Table 1 and §2.1. Subscript 1 refers to the parent, and subscript 2 refers to the offspring (the perspective from which probabilities are calculated). Circles represent living individuals, squares represent sampled individuals, and colors represent their locations: blue for the sampled parent, *x*_1_, and purple for the sampled offspring, *x*_2_. Parents are sampled on day *t*_1_, eggs are laid on day *y*_2_, and offspring are sampled on day *t*_2_. Offspring kinship probabilities are the ratio of the expected number of surviving offspring from a given adult at location *x*_2_ on day *t*_2_ (shown), and the expected number of surviving offspring from all adult females for this location and day (not shown). Calculating the expected number of surviving larval offspring at location *x*_2_ on day *t*_2_ from an adult female requires considering days of egg-laying, *y*_2_, consistent with maternal ages at sampling in the range [0, *T*_*A*_), and larval offspring ages at sampling in the range [0, *T*_*L*_). The only movement to consider is that of the mother (orange arrow).

Calculating *ψ*_*MOL*_(*x*_2_, *t*_2_ | *x*_1_, *t*_1_, *y*_2_) requires considering a single movement type - the movement of the mother between egg-laying and sampling. Given the mother is sampled at location *x*_1_ at time *t*_1_, the probability that her previous location at the time of egg-laying, *y*_2_, is *x*_2_ is calculated by normalizing over all possible egg-laying locations, i.e.:

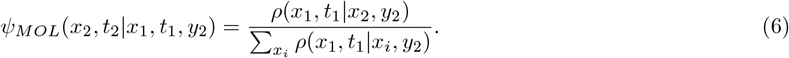

In general, *ρ*(*x*_*j*_, *t*_*j*_|*x*_*i*_, *t*_*i*_) represents the probability that an adult mosquito is at location *x*_*j*_ on day *t*_*j*_, given its location is *x*_*i*_ on day *t*_*i*_. This is given by:

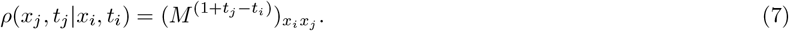

This is the (*x*_*i*_, *x*_*j*_) entry (*x*_*i*_th row and *x*_*j*_th column) of the daily movement matrix, *M*, raised to the power, (1 + *t*_*j*_ −*t*_*i*_). The power accounts for the possibility that the adult female may move on each day between egg-laying (or in later calculations, emergence) and sampling, inclusive of these two days.

Extending the mother-larval offspring kinship probability to the mother-adult offspring case is described in S1 Text §2.1. Extensions to father-offspring cases are described in S1 Text §2.2.

#### 2.2.2. Full-siblings

Next, we consider the full-sibling kinship probability for adult-adult pairs, *P*_*FSAA*_(*x*_2_, *t*_2_|*x*_1_, *t*_1_), which represents the probability that, given an adult sampled at location *x*_1_ on day *t*_1_, an adult sampled at location *x*_2_ on day *t*_2_ is their full-sibling. This can be expressed as the relative adult reproductive output at location *x*_2_ on day *t*_2_ of the mother of a larva sampled at location *x*_1_ on day *t*_1_:

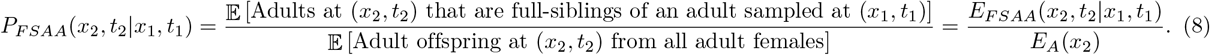

Here, *E*_*FSAA*_(*x*_2_, *t*_2_|*x*_1_, *t*_1_) represents the expected number of surviving adults at location *x*_2_ on day *t*_2_ that are full-siblings of an adult sampled at location *x*_1_ on day *t*_1_, and *E*_*A*_(*x*_2_) represents the expected number of surviving adult offspring at location *x*_2_ from all adult females at times consistent with the time of adult offspring sampling. Assuming a population at equilibrium, *E*_*A*_(*x*_2_) is independent of time and is given by:

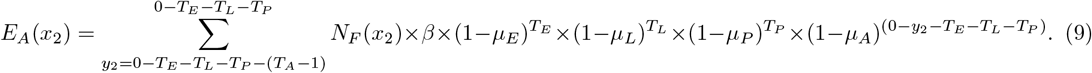

For convenience, let us refer to the adult sampled on day *t*_1_ as individual 1. To calculate *E*_*FSAA*_(*x*_2_, *t*_2_ | *x*_1_, *t*_1_), we treat the day that egg 1 is laid, *y*_1_, as a latent variable and take an expectation over it:

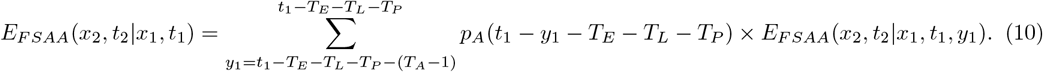

Here, the expectation over the day that egg 1 is laid, *y*_1_, is taken over days (*t*_1_ − *T*_*E*_ − *T*_*L*_ − *T*_*P*_ − (*T*_*A*_ − 1)) through (*t*_1_ − *T*_*E*_ − *T*_*L*_ − *T*_*P*_), for consistency with the day that larva 1 is sampled (Figure 3). The term *E*_*FSAA*_(*x*_2_, *t*_2_|*x*_1_, *t*_1_, *y*_1_) represents the expected number of surviving adults at location *x*_2_ on day *t*_2_ that are full-siblings of adult 1, conditional upon egg 1 being laid on day *y*_1_. Additionally, *p*_*A*_(*t*_1_ − *y*_1_ − *T*_*E*_ − *T*_*L*_ − *T*_*P*_) represents the probability that egg 1 is laid on day (*t*_1_ − *y*_1_ − *T*_*E*_ − *T*_*L*_ − *T*_*P*_). In general, *p*_*A*_(*t*) represents the probability that a given adult in the population has age *t* which, following from the daily adult survival probability, (1 − *µ*_*A*_), is given by:

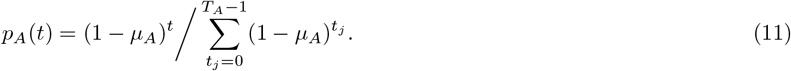

**Fig 3.**
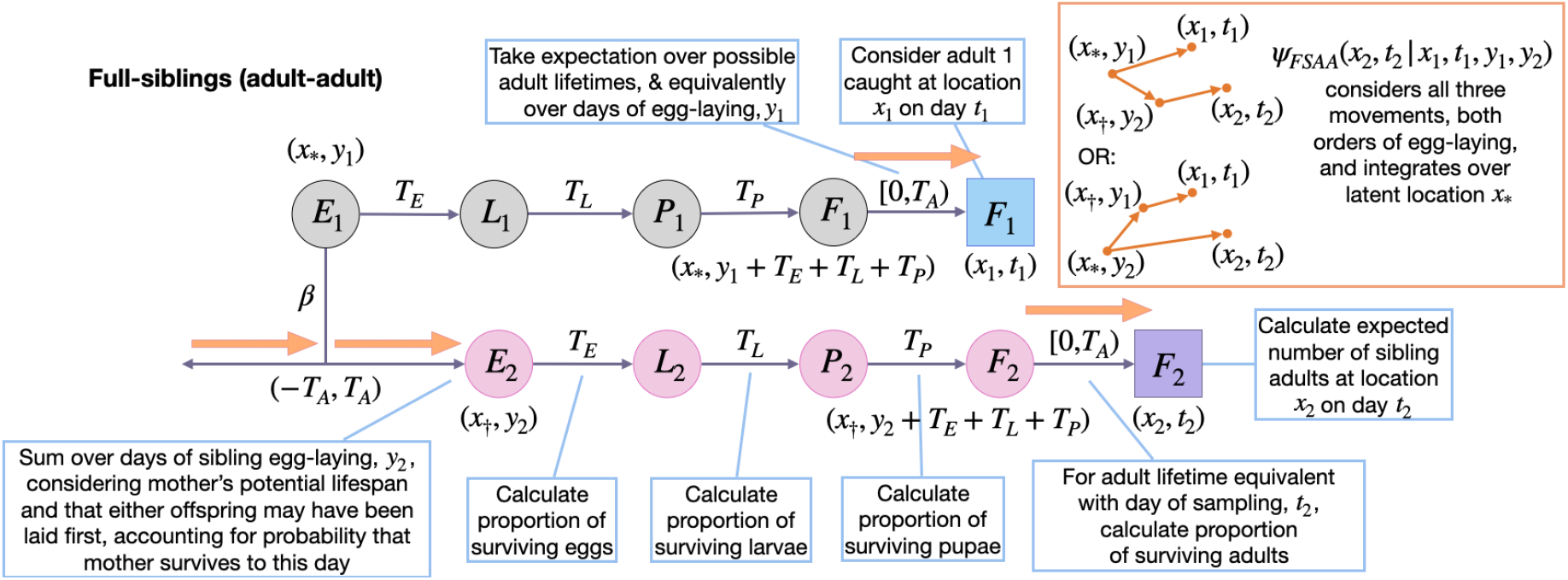
Schematic representation of spatial adult-adult full-sibling kinship probabilities. Parameters and state variables are as defined in Table 1 and §2.2. Subscript 1 refers to the reference sibling, and subscript 2 refers to the sibling from whose perspective the probabilities are calculated. Circles represent living individuals, squares represent sampled individuals, and colors represent their locations: blue for sibling 1, *x*_1_, purple for sibling 2, *x*_2_, grey for the location of egg-laying for sibling 1, and pink for the location of egg-laying for sibling 2. The location of egg-laying for the firstborn sibling (here, sibling 1) is denoted by *x*_*\**_, and *x*_*†*_ denotes the egg-laying location for the other sibling. Sibling 1 is sampled on day *t*_1_ and laid on day *y*_1_. Sibling 2 is sampled on day *t*_2_ and laid on day *y*_2_. Sibling kinship probabilities are the ratio of the expected number of surviving siblings of a given individual at location *x*_2_ on day *t*_2_ (shown), and the expected number of surviving offspring from all adult females for this location and day (not shown). Calculating the expected number of surviving full-siblings at location *x*_2_ on day *t*_2_ requires considering days of egg-laying, *y*_1_ and *y*_2_, consistent with adult ages at sampling in the range [0, *T*_*A*_). There are three movements to consider: those of the mother and two adult siblings (orange arrows). Movement probabilities consider both orders of egg-laying.

The term *E*_*FSAA*_(*x*_2_, *t*_2_|*x*_1_, *t*_1_, *y*_1_) represents the expected number of surviving adults at location *x*_2_ on day *t*_2_ that are full-siblings of adult 1, conditional upon egg 1 being laid on day *y*_1_. This is given by:

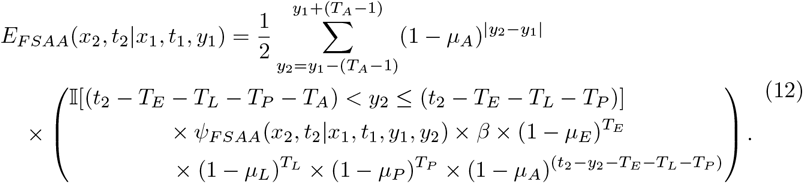

Here, the day of sibling egg-laying, *y*_2_, is summed over days (*y*_1_ − (*T*_*A*_ − 1)) through (*y*_1_ + (*T*_*A*_ − 1)), for consistency with the mother’s potential lifespan (Figure 3), and considering that either of the larval offspring may have been laid first. Since we are summing two equally-weighted scenarios regarding offspring order, we include a multiplier of 1*/*2 in the expectation. The first term within the summation then represents the probability that the mother is alive on the day of sibling egg-laying, with the absolute value, |*y*_2_ − *y*_1_|, accounting for both offspring orders. The second term (in larger brackets) represents the expected adult offspring output of the mother of adult 1 at location *x*_2_ on day *t*_2_. This is the same equation as for the mother-larval offspring case with three exceptions: i) daily egg production is multiplied by the proportion of eggs that survive the egg, larva, pupa and adult stages up to the day of sampling to reflect the fact that adults rather than larvae are being sampled, ii) the indicator function limits consideration to cases where the day of sibling egg-laying, *y*_2_, is between days (*t*_2_ − *T*_*E*_ − *T*_*L*_ − *T*_*P*_ − (*T*_*A*_ − 1)) and (*t*_2_ − *T*_*E*_ − *T*_*L*_ − *T*_*P*_), for consistency with an adult sibling being sampled on day *t*_2_, and iii) there are now three movements captured by the composite movement term, *ψ*_*FSAA*_(*x*_2_, *t*_2_|*x*_1_, *t*_1_, *y*_1_, *y*_2_), which represents the probability that adult offspring 2 is sampled at location *x*_2_ on day *t*_2_ given that adult offspring 1 is sampled at location *x*_1_ on day *t*_1_, egg 1 is laid on day *y*_1_, and egg 2 is laid on day *y*_2_.

Calculating *ψ*_*FSAA*_(*x*_2_, *t*_2_ | *x*_1_, *t*_1_, *y*_1_, *y*_2_) requires considering the mother’s movement between egg-laying events, in addition to the movement of both adult offspring for both offspring orders, i.e.:

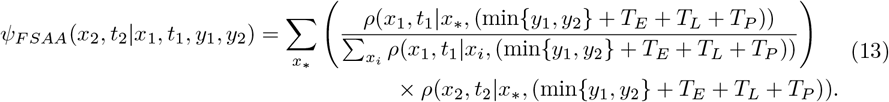

Here, we take an expectation over a latent egg-laying location for the firstborn sibling, *x*_*\**_, and multiply the probability that the firstborn sibling is laid at location *x*_*\**_ given that adult sibling 1 is sampled at location *x*_1_ by the probability that adult sibling 2 is sampled at location *x*_2_ (the former probability requires normalizing over all egg-laying locations). For both adult siblings, movement begins after development through the egg, larva and pupa life stages (i.e., after (*T*_*E*_ + *T*_*L*_ + *T*_*P*_) days), and the mother’s movement between egg-laying events is incorporated by effectively adding |*y*_2_ − *y*_1_| days of movement either backwards or forwards in time, depending on the offspring order.

Extending full-sibling kinship probabilities to other life stage pairs is relatively straightforward. We consider the case of larva-larva, larva-adult and adult-larva full-sibling pairs in S1 Text §2.3.

### 2.3. Pseudo-likelihood calculation

The goal of this spatial CKMR analysis is to make inferences about dispersal parameters given data on the frequency, timing and location of observed close-kin pairs.

Here, we calculate the likelihood of parent-offspring and full-sibling pairs in a manner that takes advantage of the nature of the kinship probabilities and the sampling process. The kinship probabilities for each pair of individuals are assumed to be independent of each other, even though they are not. For this reason the combined likelihood is referred to as a “pseudo-likelihood” [9] - an approach that has been shown to produce accurate parameter and variance estimates provided the size of each sampling event is sufficiently low relative to the total population size [28, 29].

#### 2.3.1. Parent-offspring pairs

Let us begin by considering the mother-larval offspring kinship probability, *p*_*MOL*_(*x*_2_, *t*_2_|*x*_1_, *t*_1_), which represents the probability that, given an adult female sampled at location *x*_1_ on day *t*_1_, a given larva sampled at location *x*_2_ on day *t*_2_ is her offspring. Now consider *n*_*F*_ (*x*_1_, *t*_1_) adult females sampled at location *x*_1_ on day *t*_1_. The probability that a larva sampled at location *x*_2_ on day *t*_2_ has a mother amongst the *n*_*F*_ (*x*_1_, *t*_1_) sampled adult females, *p*_*MOL*_(*x*_2_, *t*_2_|*x*_1_, *t*_1_), is equal to one minus the probability that none of the *n*_*F*_ (*x*_1_, *t*_1_) sampled adult females are the larva’s mother, i.e.:

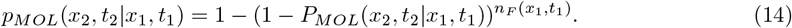

Here, *P*_*MOL*_(*x*_2_, *t*_2_|*x*_1_, *t*_1_) is as defined in Equation 3. Now consider *n*_*L*_(*x*_2_, *t*_2_) larvae sampled at location *x*_2_ on day *t*_2_, and let *k*_*MOL*_(*x*_2_, *t*_2_|*x*_1_, *t*_1_) be the number of larvae sampled at location *x*_2_ on day *t*_2_ that have a mother amongst the adult females sampled at location *x*_1_ on day *t*_1_. The pseudo-likelihood that *k*_*MOL*_(*x*_2_, *t*_2_|*x*_1_, *t*_1_) of the *n*_*L*_(*x*_2_, *t*_2_) larvae sampled at location *x*_2_ on day *t*_2_ have a mother amongst the adult females sampled at location *x*_1_ on day *t*_1_ follows from the binomial distribution:

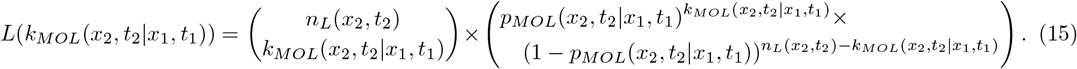

The full log-pseudo-likelihood for mother-larval offspring pairs, Λ_*MOL*_, follows from summing the log-pseudo-likelihood over all adult female sampling days, *t*_1_, over consistent larval offspring sampling days, *t*_2_, and over all adult female and larval sampling locations, *x*_1_ and *x*_2_, respectively:

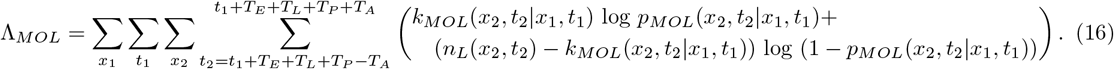

Note that, for the purpose of parameter interference, we can drop the first term in the pseudo-likelihood equation, and for the purpose of efficient computation, we consider consistent adult sampling days from (*t*_1_ + *T*_*E*_ + *T*_*L*_ + *T*_*P*_ − (*T*_*A*_ − 1)) through (*t*_1_ + *T*_*E*_ + *T*_*L*_ + *T*_*P*_ + (*T*_*A*_ − 1)). The earliest adult sampling day (relative to *t*_1_) corresponds to the case where the mother laid the offspring at the beginning of her life, was sampled at the end of her life, and the adult offspring was sampled at the beginning of its life. The latest adult sampling day (relative to *t*_1_) corresponds to the case where the mother was sampled on the day they laid their offspring, and the adult offspring was sampled at the end of its life.

Parent-offspring pseudo-likelihood equations for other sampled sexes and life stages follow an equivalent formulation. The main point to note is that consistent offspring sampling days are specific to the kinship and sampled life stages being considered (these can be deduced from event history diagrams like those in Figure 2). For adult offspring cases where *x*_1_ = *x*_2_ and *t*_1_ = *t*_2_, the number of sampled adults, *n*_*A*_(*x*_2_, *t*_2_), is reduced by one to account for the fact that an adult cannot be its own parent. The joint log-pseudo-likelihood for parent-offspring pairs is then given by:

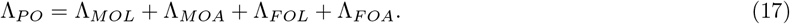

Here, Λ_*MOA*_, Λ_*FOL*_ and Λ_*FOA*_ denote the log-pseudo-likelihoods for mother-adult offspring pairs, father-larval offspring pairs and father-adult offspring pairs, respectively.

#### 2.3.2. Full-sibling pairs

For full-siblings, we begin with the adult-adult full-sibling kinship probability, *P*_*FSAA*_(*x*_2_, *t*_2_|*x*_1_, *t*_1_), defined in Equation 8, which represents the probability that, given an adult sampled at location *x*_1_ on day *t*_1_, an adult sampled at location *x*_2_ on day *t*_2_ is their full-sibling. We consider a given adult, indexed by *i* and sampled at location *x*_1_(*i*) on day *t*_1_(*i*), and *n*_*A*_(*x*_2_, *t*_2_) adults sampled at location *x*_2_ on day *t*_2_. Let *k*_*FSAA*_(*i, x*_2_, *t*_2_) be the number of adults sampled at location *x*_2_ on day *t*_2_ that are full-siblings of adult *i*. The pseudo-likelihood that *k*_*FSAA*_(*i, x*_2_, *t*_2_) of the *n*_*A*_(*x*_2_, *t*_2_) sampled adults at location *x*_2_ on day *t*_2_ are full-siblings of adult *i* follows from the binomial distribution:

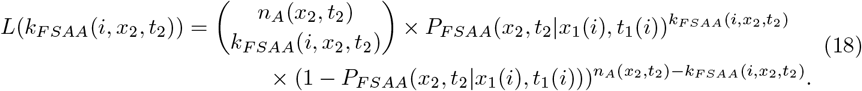

Note that, for cases where *x*_1_(*i*) = *x*_2_ and *t*_1_(*i*) = *t*_2_, the number of sampled adults at location *x*_2_ on day *t*_2_, *n*_*A*_(*x*_2_, *t*_2_), is reduced by one to account for the fact that an adult cannot be its own sibling. Additionally, when counting siblings, we only consider siblings with indices > *i* to avoid double-counting. The full log-pseudo-likelihood for adult-adult full-sibling pairs, Λ_*FSAA*_, follows from summing the log-pseudo-likelihood over all sampled adults, *i*, over all sampled locations, *x*_2_, and over consistent adult sampling days, *t*_2_:

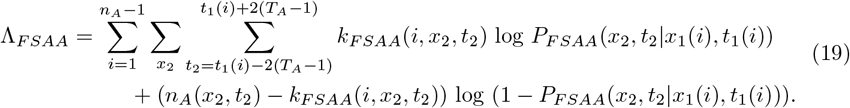

Consistent adult sampling days for this case are from (*t*_1_(*i*) − 2(*T*_*A*_ − 1)) through (*t*_1_(*i*) + 2(*T*_*A*_ − 1)). The earliest adult sampling day (relative to *t*_1_(*i*)) corresponds to the case where the mother laid individual 2 at the beginning of her life and individual 1 at the end of her life, adult 1 was sampled at the end of its life, and adult 2 was sampled soon after emergence. The latest adult sampling day (relative to *t*_1_(*i*)) corresponds to the reverse case. Full-sibling pseudo-likelihood equations for other life stage pairs follow an equivalent formulation, with consistent sampling days specific to the kinship and sampled life stages being considered (these can be deduced from event history diagrams like those in Figure 3). The joint log-pseudo-likelihood for full-sibling pairs is then given by:

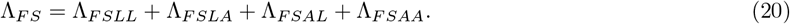

Here, Λ_*FSLL*_, Λ_*FSLA*_ and Λ_*FSAL*_ denote the log-pseudo-likelihoods for larva-larva, larva-adult and adult-larva full-sibling pairs, respectively.

#### 2.3.4. Parameter inference

Despite parent-offspring and full-sibling kinship probabilities not being independent, the pseudo-likelihood approach enables us to combine these likelihoods, provided the size of each sampling event is sufficiently low relative to the total population size [9]. As we will see later, our simulation studies suggest this to be the case. We therefore combine these log-pseudo-likelihoods to obtain a log-pseudo-likelihood for the entire data set:

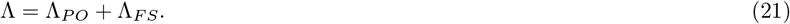

Parameter inference can then proceed by varying a subset of the dispersal and/or demographic parameters in Table 1 in order to minimize −Λ. We used the **nlminb** function implemented in the **optimx** function in R [30] to perform our optimizations.

### 2.4. Individual-based simulation model

We used a previously-developed simulation package, **mPlex** [16], to model mosquito life history and to test the effectiveness of the CKMR approach at estimating mosquito dispersal and demographic parameters. The model is an individual-based adaptation of a previous model, **MGDrivE** [31], which is a genetic and spatial extension of the lumped age-class model applied to mosquitoes by Hancock and Godfray [19] and Deredec *et al*. [20] (Figure 1A). Our previous application of **mPlex** to CKMR-based inference problems considered only panmmictic populations; however, the functionality had already been included to account for spatial population structure, with mosquitoes being distributed across populations in a metapopulation [31]. Each population has an equilibrium adult population size, 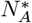, and exchanges migrants with the other populations. Populations are partitioned according to discrete life stages - egg, larva, pupa and adult - with sub-adult stages having fixed durations as defined earlier. See Sharma *et al*. [16] for more details.

## 3. Results

We used simulated data from the individual-based mosquito model to explore the feasibility of spatial CKMR to infer dispersal parameters for *Ae. aegypti*. Our simulated metapopulation consisted of a 19-by-19 grid of households (Figure 1C) each inhabited by 25 adult mosquitoes at equilibrium with bionomic parameters listed in Table 1. Landscape dimensions were chosen to accommodate the trap arrangements described below, as well as a buffer width of at least three non-trap nodes along each landscape edge (e.g., Figure 1D) to reduce boundary effects. Open questions concern the optimal sampling scheme to estimate dispersal parameters for *Ae. aegypti* using spatial CKMR methods, and the range of dispersal parameters that can be accurately estimated. To address these questions, we first explored logistically-feasible sampling schemes to estimate mean dispersal distance by varying: i) sampled life stage (larva or adult), ii) total sample size (1,000-3,000 sequenced individuals), and iii) the number and spacing of trap nodes (arranged in 4-by-4, 5-by-5 or 6-by-6 grids with zero, one or two population nodes separating each trap node). Based on a previous analysis to estimate mosquito demographic parameters using CKMR [16], our adult samples were of females (since mosquito traps are often tailored to this sex), sampling frequency was biweekly (as is often the case for mosquito surveillance programs [32]), and sampling duration was for three months (corresponding to a season). Our likelihood calculations were based on parent-offspring and full-sibling pairs. Half-sibling pairs could be included for individual data analyses; however, computational burden prevented us from including these for exploratory analyses. Initially, we focused on estimating mean daily dispersal distance, 1*/λ*_*d*_, and for subsequent analyses, also estimated daily staying probability, *p*_0_, and barrier strength, *δ*.

### 3.1. Optimal sampling scheme to estimate daily dispersal distance

To estimate mean daily dispersal distance, our default sampling scheme consisted of a total of 1,000 sequenced individuals sampled biweekly over a three-month period spread over a 6-by-6 grid of trap nodes with one population node separating each trap node (Figure 1C) (i.e., ca. 1 individual sampled twice per week from each trap node, for a total of 1,000 individuals across all trap nodes after three months of sampling). We first explored the optimal distribution of sampled life stage to estimate 1*/λ*_*d*_, exploring three scenarios: all larvae, all adult females, and half larvae/half adult females. Results of 100 simulation-and-analysis replicates for each scenario are depicted in Figure 4A. These suggest that all three life stage scenarios result in adequate parameter inference, in terms of both accuracy of the median and tightness of the interquartile range (IQR), albeit consistently underestimating 1*/λ*_*d*_ by ca. 10-15%. Since most mosquito traps are designed to target adult females, we will proceed with modeling samples of this life stage. While larval samples also produce adequate inference, either exclusively or together with adult female samples, larvae are more difficult to sample in the environment due to breeding sites sometimes being hidden or inaccessible. The problem of larval sampling is exacerbated for CKMR studies due to the requirement that individuals be sampled independently. Therefore, if multiple larvae are collected in a larval dip from a single breeding site, only one can be used in the analysis to prevent biasing the number of collected sibling pairs upwards, thus further increasing the effort required to achieve a substantial larval sample.

**Fig 4.**
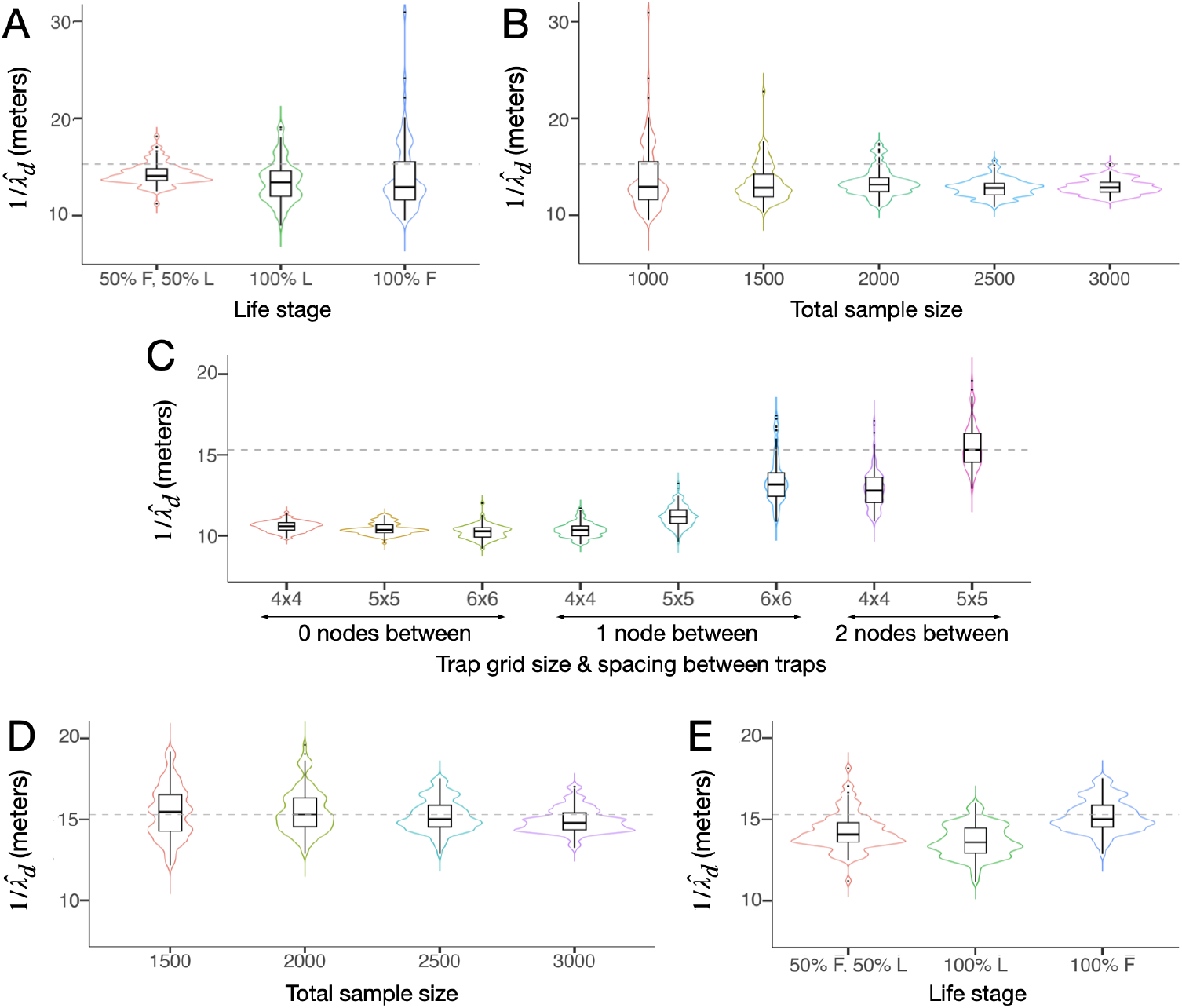
Sampling schemes to estimate 1/*λ*_*d*_ for *Ae. aegypti*. Violin plots depict estimates of 1/*λ*_*d*_ for sampling scenarios described in §3.1. The simulated metapopulation consists of a 19-by-19 grid of households each inhabited by 25 adult *Ae. aegypti* at equilibrium with bionomic parameters listed in Table 1. Boxes depict median and interquartile ranges of 100 simulation-and-analysis replicates for each scenario, thin lines represent 5% and 95% quantiles, points represent outliers, and kernel density plots are superimposed. The default sampling scheme consists of a total of 1,000 individuals sampled as ca. 1 individual collected twice weekly over a three-month period for each trap node, considering a 6-by-6 array of trap nodes with one population node separating each trap node (Figure 1C). In panel **(A)**, three life stage proportions are explored: all larvae, all adult females, and half larvae/half adult females. In panel **(B)**, all sampled individuals are adult females, and total sample sizes of 1,000, 1,500, 2,000, 2,500 and 3,000 are explored. In panel **(C)**, a sample size of 2,000 is adopted, and the number and spacing of trap nodes is varied (arranged in 4-by-4, 5-by-5 or 6-by-6 grids with zero, one or two population nodes separating each trap node). In panel **(D)**, trap nodes are arranged in a 5-by-5 grid with two population nodes separating each trap node (Figure 1D), and the optimal total sample size is revisited. In panel **(E)**, a sample size of 2,500 adult females is adopted, and the sampled life stage is revisited. The optimal sampling scheme consists of 2,500 adult females collected biweekly over a three-month period spread over a 5-by-5 grid of trap nodes with two population nodes separating each trap node.

Next, we explored the optimal sample size to estimate 1/*λ*_*d*_ for *Ae. aegypti*. We performed 100 simulation-and-analysis replicates for each of five total sample sizes - 1,000, 1,500, 2,000, 2,500 and 3,000 adult females - depicted in Figure 4B. Results suggest that estimates of 1/*λ*_*d*_ become more precise for larger sample sizes (as measured by the IQR); but there are diminishing returns in precision for sample sizes larger than 2,000. The median estimate of 1/*λ*_*d*_ remains underestimated by ca. 10-15% regardless of sample size. We therefore proceed with an optimal sample size of 2,000 adult females, collected biweekly over a three-month period. This equates to ca. 2 individuals sampled twice per week from each trap node over the three-month period.

Next, we explored the optimal number and spacing of trap nodes to estimate 1/*λ*_*d*_ for *Ae. aegypti*. We performed 100 simulation-and-analysis replicates for trap nodes arranged in 4-by-4, 5-by-5 or 6-by-6 grids with zero, one or two population nodes separating each trap node (except for the case of a 6-by-6 grid of trap nodes where the simulated landscape could only accommodate zero or one population node separating each trap node). Results, depicted in Figure 4C, suggest that when traps are optimally spread out throughout a landscape, inference of mean dispersal distance is very accurate (as measured by the difference between the median and true value). Notably, estimates of 1*/λ*_*d*_ are unbiased for trap nodes arranged in a 5-by-5 grid with two population nodes separating each trap node, and are relatively close to the true value for a smaller number of trap nodes (e.g., arranged in a 4-by-4 grid with two population nodes separating each trap node), or for a larger number of trap nodes that are closer together (e.g., arranged in a 6-by-6 grid with one population node separating each trap node). Conversely, estimates of 1*/λ*_*d*_ are highly biased when traps are clustered together, as can be seen for grids of 4-by-4, 5-by-5 or 6-by-6 adjacent trap nodes. In these cases, the median estimate of 1*/λ*_*d*_ is ca. 30% less than the true value. This is an intuitive result, as when trap nodes are clustered together, they are more likely to capture nearby movement but less likely to capture distant movement, hence leading to underestimates of mean daily dispersal. The optimal sampling scenario where trap nodes are separated by two population nodes equates to a distance between trap nodes of 49.8 meters (since population nodes are separated by 16.6 meters in these simulations), which is approximately equal to the mean lifetime dispersal distance of *Ae. aegypti* mosquitoes used to parameterize this model of 45.2 meters [11]. Finally, the 5-by-5 grid of trap nodes with each trap node separated by two population nodes represents the largest number of traps that can be accommodated on the simulated landscape for this degree of separation.

Finally, given the significance of the number and spacing of trap nodes for estimating 1/*λ*_*d*_ evident in Figure 4C, we revisited the optimal sample size and life stage distribution given a 5-by-5 grid of trap nodes with each trap node separated by two population nodes. Results for sample size again suggest that samples of 2,000 adult females produce adequate estimates of 1/*λ*_*d*_; but also suggest an improvement in precision (as measured by IQR) for a population size of 2,500 (Figure 4D). We therefore proceed with this total sample size, which equates to ca. 3-4 individuals sampled twice per week for each trap node over the three-month period. Regarding the optimal distribution of sampled life stage, results again suggest that all three life stage scenarios result in comparable parameter inference (as measured by the accuracy of the mean and the tightness of the IQR) (Figure 4E). We therefore continue with an adult female sample, as per our previous reasoning. With all of these considerations in mind, the optimal sampling scheme therefore consists of a total of 2,500 adult females collected biweekly over a three-month period spread over a 5-by-5 grid of trap nodes with two population nodes separating each trap node.

### 3.2. CKMR-based estimates of barrier strength and daily staying probability

Given the optimal sampling scheme, we employ the flexibility of the formal CKMR approach to estimate additional parameters describing more complex dispersal patterns - the strength of a barrier to movement, *δ*, and the daily staying probability for a zero-inflated exponential kernel, *p*_0_. To begin, we consider a barrier to movement, which could represent a road, freeway or open park space for *Ae. aegypti*, as has been documented as being important for dispersal of this species [21]. We depict the barrier as a line through the landscape (Figure 1D), whereby movement to the other side of the barrier is reduced by a factor, *δ*, and movement on the same side of the barrier is unaltered. We explore *δ* values in the range [0.1, 0.9] and adopt the optimal sampling scheme determined in §3.1 - a total sample of 2,500 adult females sampled from a 5-by-5 grid of trap nodes with two population nodes separating each trap node. Results in Figure 5C suggest that estimates of *δ* are very accurate (as measured by the distance between the median of 100 simulations and the true value) for *δ* ≥ 0.5, and barrier strength estimates become more precise (as measured by IQR) for *δ* ≥ 0.75. Estimates of barrier strength tend to be slightly overestimated for *δ ≤* 0.25; but the true value still falls within the IQR. Estimates of 1/*λ*_*d*_ are reasonably accurate when barrier strength is simultaneously estimated (Figure 5A), with the true value consistently falling within the IQR; however, the median estimate from 100 simulations falls below the true value by ca. 3-5% for barrier strengths ≥ 0.75. This is perhaps not surprising, considering that the presence of a strong barrier in a landscape reduces the mean distance traveled by an individual over their lifetime.

**Fig 5.**
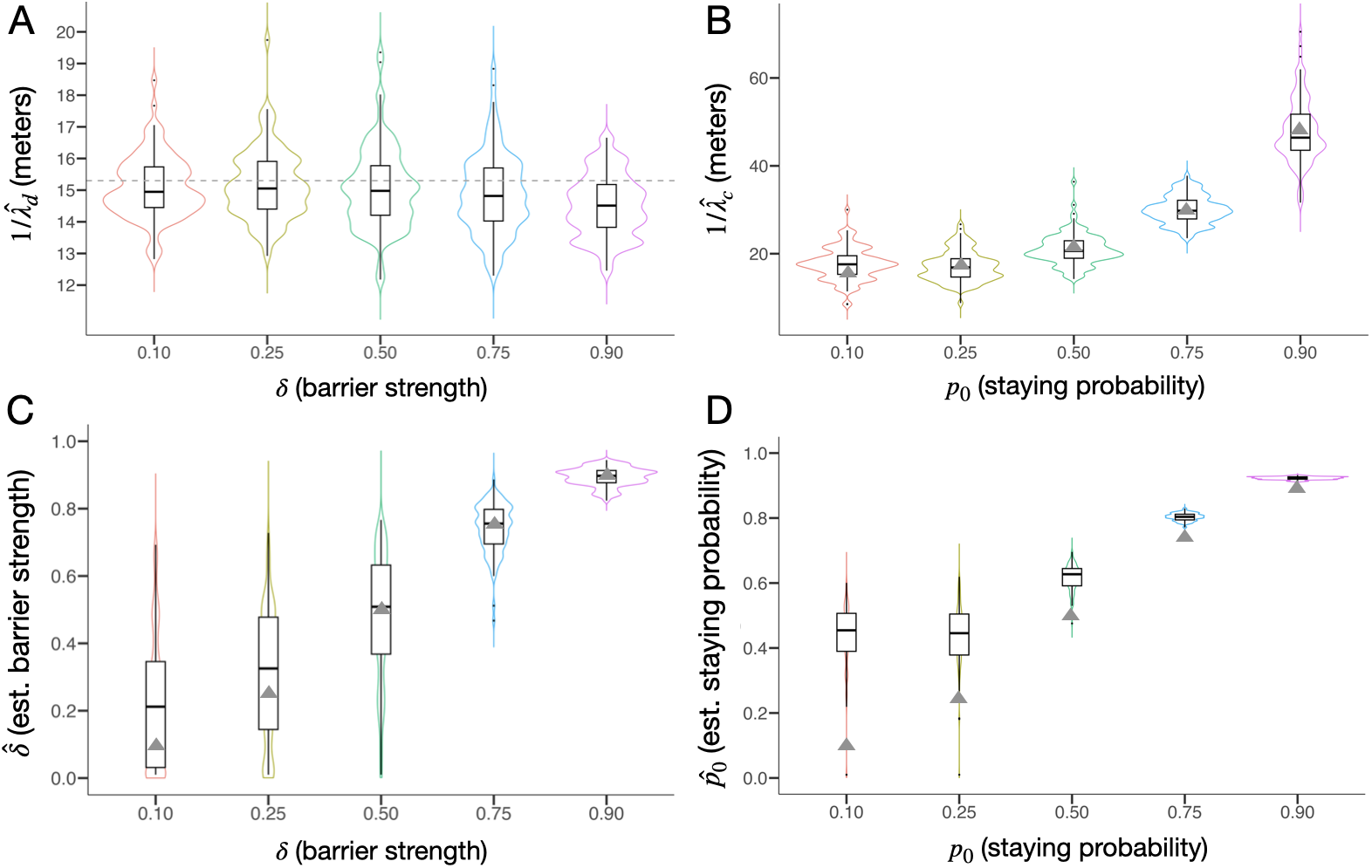
Estimates of barrier strength and daily staying probability using spatial CKMR methods. In the first (left) analysis, violin plots depict estimates of mean daily dispersal distance, 1*/λ*_*d*_ **(A)**, and barrier strength, *δ* **(C)**, obtained using the spatial CKMR approach for the optimal sampling scheme determined in §3.1, and considering a barrier to movement as depicted in Figure 1D whereby movement to the other side of the barrier is reduced by a factor, *δ*, in the range [0, 0.9]. In the second (right) analysis, violin plots depict estimates of mean daily dispersal distance conditional upon movement, 1/*λ*_*c*_ **(B)**, and daily staying probability, *p*_0_ **(D)**, in the range [0.1, 0.9], again obtained using the spatial CKMR approach for the optimal sampling scheme determined in §3.1, and considering a zero-inflated dispersal kernel as described in Equation 2 and depicted in Figure 1B. The simulated metapopulation consists of a 19-by-19 grid of households each inhabited by 25 adult *Ae. aegypti* at equilibrium with bionomic parameters listed in Table 1. Boxes depict median and interquartile ranges of 100 simulation-and-analysis replicates for each scenario, thin lines represent 5% and 95% quantiles, points represent outliers, and kernel density plots are superimposed. True parameter values are depicted by triangles, or by a dotted line when they are consistent across all cases.

Next, we consider a zero-inflated exponential dispersal kernel and estimate the daily staying probability, *p*_0_, alongside the mean daily dispersal distance conditional upon movement, *λ*_*c*_. Exploring this kernel is motivated by the fact that several mosquito species, such as *Ae. aegypti*, obtain most of their resources from a small area such as a household and tend to remain at this location for the majority of their lifetime [5]. The zero-inflated exponential dispersal kernel is formulated in Equation 2, and dispersal kernels having *p*_0_ ∈ [0.1, 0.9] are depicted in Figure 1B. For each value of *p*_0_, we maintain a mean daily dispersal distance of 15.3 m (Table 1), which translates to a daily dispersal distance conditional upon movement, 1/*λ*_*c*_, of 16.1 m for *p*_0_ = 0.1, 17.7 m for *p*_0_ = 0.25, 21.6 m for *p*_0_ = 0.5, 30.6 m for *p*_0_ = 0.75, and 48.4 m for *p*_0_ = 0.9. Again, we assume the optimal sampling scheme determined in §3.1. Results in Figure 5B suggest that estimates of 1/*λ*_*c*_ are accurate and precise for all explored values of *p*_0_, as measured by the difference between the median of 100 simulations and the true value of 1/*λ*_*c*_, and by the IQR. On the other hand, *p*_0_ can be accurately and precisely estimated for true *p*_0_ ≥ 0.75; but smaller true values of *p*_0_ are overestimated by increasingly large degrees (as measured by the median of 100 simulation replicates) - a true value of *p*_0_ of 0.5 is overestimated by ca. 0.09, a true value of 0.25 is overestimated by ca. 0.20, and a true value of 0.1 is overestimated by ca. 0.36 (Figure 5D). Overestimates of *p*_0_ may be related to the calculation of movement probabilities on a grid landscape. For the 19-by-19 landscape depicted in Figure 1C, an exponential kernel without zero-inflation has diagonal entries of its transition matrix (effectively, staying probabilities) between 0.18 and 0.38, and hence staying probabilities of a zero-inflated kernel with *p*_0_ ∈ {0.1, 0.25} may be difficult to discern.

## 4. Discussion

We have demonstrated that the CKMR formalism with spatial structure described by Bravington *et al*. [9] can be used to estimate dispersal parameters of *Ae. aegypti*, a major vector of arboviruses such as dengue and Zika virus, as a case study. Using a spatial individual-based simulation framework [16] based on the lumped age-class model [17] applied to mosquitoes [19], we have shown that these methods accurately estimate mean daily dispersal distance, 1*/λ*_*d*_, and can also estimate parameters of more complicated dispersal kernels, such as the strength of a barrier to movement, *δ*, and the daily staying probability, *p*_0_, for a zero-inflated exponential kernel. The optimal sampling scheme inferred in this study is consistent with standard *Ae. aegypti* field sampling protocols, provided the distribution of traps is carefully selected. As for a previous *in silico* mosquito CKMR analysis [16], we found that sampling adult females biweekly over a period of three months is adequate, as is commonplace for mosquito sampling [32] and consistent with the length of a season.

The simulated 19-by-19 grid landscape with nodes (in this case, households) spaced 16.6 meters apart was designed to resemble a suburban setting such as that of Cairns, Australia, as a case study (the neighborhood dimensions were chosen to resemble those of the Cairns suburb, Yorkeys Knob). In this setting, the optimal explored sampling scheme consisted of a five-by-five grid of trap nodes with two household nodes separating each trap node. Of note, traps in this configuration were separated by approximately the simulated lifetime dispersal distance (i.e., a separation distance of 49.8 meters c.f. a lifetime dispersal distance of 45.2 meters [11]), with the largest number of traps that could be accommodated on the simulated landscape being optimal. This optimal trap layout and the optimal sample size of 2,500 adult females is amenable to *Ae. aegypti* field protocols. Dividing this total sample size across the full network of traps and a three-month sampling period equates to ca. 3-4 adult females sampled twice-weekly for each trap node. This sample size is achievable; but were it to present a challenge, then additional traps could be placed throughout the landscape. It is worth noting that smaller sample sizes would likely be required for smaller populations (our simulations assumed 25 adult mosquitoes per household [22]).

This formal CKMR approach to estimating mosquito dispersal parameters is complementary to the recently-proposed methods of Filipović *et al*. [11] and Jasper *et al*. [10], with each method having its own strengths and weaknesses. The Filipović *et al*. method is described in full in the Methods section of [11]; but in brief, adult females are captured while ovipositing, kinship categories determined, and a mean generational displacement calculated for each close-kin pair. This calculation considers the accumulated displacement between the close-kin individuals, and the set of possible movement events that led to it. A dispersal kernel is fitted to the set of mean generational displacements for all close-kin pairs. The Jasper *et al*. method is described in full in the Materials and Methods section of [10]; but in brief, eggs are collected from ovitraps, kinship categories determined, and “axial standard deviations” are calculated for each kinship category. The mean dispersal distance of adult females between emergence and egg-laying is then calculated from variance formulae that incorporate the axial standard deviations of observed kinship categories (see Equations 1-3 of [10]). This method can also be applied to sampled adult females, as demonstrated in [33], although it performs better when applied to sampled eggs.

Key strengths of the Jasper *et al*. [10] and Filipović *et al*. [11] methods are that: i) they are simpler than the formal CKMR approach, and hence require less computational investment, and ii) they accommodate second and third-degree close-kin without computational burden. This however makes a systematic comparison of the performance of the three methods difficult, as the Jasper *et al*. and Filipović *et al*. methods are ideally performed on larger landscapes that accommodate displacement accumulated over 2-3 generations, while the formal CKMR approach can be implemented on a smaller landscape, inferring mean daily dispersal from displacement accumulated over shorter time periods spanning one day through two generations. Of note, the Filipović *et al*. method can be applied exclusively to first-degree close-kin, making a reduced landscape amenable to that approach; however, this comes at the expense of sacrificing second and third-degree close-kin data that the method otherwise benefits from. For the formal CKMR approach, the incorporation of temporal information enables higher-resolution inference of dispersal parameters from data collected over a smaller area; however, the computational burden associated with this additional detail limits the size of a landscape that can be considered using this method.

A key benefit of the formal CKMR approach is its ability to estimate parameters of more complex dispersal kernels, such as the staying probability of a zero-inflated exponential kernel, and of more complex landscapes, such as the strength of a barrier to dispersal. Exponential and zero-inflated exponential kernels were explored in this analysis; but any number of dispersal kernels could be explored, provided available data is consistent with identifiability of their parameters. A related benefit of the CKMR approach is that it can be tailored to the life history and landscape of a specific mosquito species and location. This includes three-dimensional landscapes such as the multi-storey housing blocks analyzed in Singapore by Filipović *et al*. [11]. It should be noted that, with this capability comes a need to understand the local ecology of the species before parameter inference begins. E.g., when estimating dispersal parameters in this study, we incorporated a fully-specified life history model in addition to knowledge of the distribution of habitat patches and the functional form of the dispersal kernel. That said; the formal CKMR approach can also be used to estimate demographic parameters unrelated to dispersal, such as census population size, adult and larval daily mortality rates, and larval life stage duration, as shown in Sharma *et al*. [16].

As a preliminary exploration of the application of formal CKMR methods to estimate dispersal parameters of mosquitoes, this study has several limitations. First, the same life history and landscape model (Figure 1) was used as a basis for both the population simulations and CKMR analysis. Additionally, other than the parameters being estimated, the same parameters were used in both the simulations and analysis. This represents an overly generous scenario as compared to the field, where life history is varied and complex and parameters are only approximately known. That said; this is an appropriate starting point to verify the utility of the method. Second, we have assumed perfect kinship inference throughout. Incorporating kinship uncertainty into the CKMR likelihood equations is theoretically possible [34], although this has produced little improvement in parameter inference at large computational cost when applied to data from fish species [28]. One approach to assess the importance of kinship assignment errors would be to introduce them at the simulation phase, and to then test the robustness of the methods to these errors. Here, there is an important distinction between type I (false positive) and II (false negative) error rates. Studies in fish species suggest that kinship inference must have a very low type I error rate in order for CKMR parameter inference to be informative [9].

The application of formal CKMR methods to spatial settings, while contemplated since their inception [9], has only been considered in a small number of cases for species with simpler life histories [14, 15]. The extension of these methods to a metapopulation of insects having an egg-larva-pupa-adult life history is promising for insights this approach may provide for other species, alongside insights from simpler close-kin-based approaches [10, 11]. Potential applications to *An. gambiae* are of particular interest, given the importance of this species for malaria transmission and the importance of quantifying dispersal patterns for planning vector control interventions and field trials. The increased dispersal range of this species is important to acknowledge [5], as is the potential for age-grading techniques [35] to enhance parameter inference. Several species of insect agricultural pests share a similar life history, and close-kin-based approaches such as CKMR should be explored to provide insight into their dispersal patterns.

## 5. Conclusions

We have theoretically demonstrated the application of spatial CKMR methods to estimate dispersal parameters for mosquitoes, with *Ae. aegypti* as a case study. Close-kin-based methods have advantages over traditional MRR methods, as the mark is genetic, removing the need for physical marking and recapturing. Encouragingly, we find that optimal spatial CKMR sampling schemes are consistent with *Ae. aegypti* ecology and field studies, provided the spatial distribution of traps is carefully chosen. In our *in silico* case study, we found that traps distributed in a grid layout separated at a distance approximately equal to the mean lifetime dispersal distance of this species were optimal. The formal CKMR approach is complementary to two simpler close-kin-based methods at estimating mean dispersal distance, with each approach having its own strengths and weaknesses. The CKMR approach is particularly computationally intensive, restricting its ability to be applied to second and third-degree close-kin and to larger landscapes; but its ability to be tailored to a specific landscape and dispersal kernel enable it to estimate additional parameters such as the daily staying probability and strength of a barrier to movement. Close-kin-based methods such as CKMR promise to provide further insight into the dispersal patterns of other insects of epidemiological and agricultural significance.

## Supporting information

**S1 Text. Supplemental model equations**. Additional equations describing mosquito dispersal dynamics, and spatial kinship probabilities for parent-offspring and full-sibling pairs that, for brevity, were not included in the manuscript.

## Acknowledgments

We thank Dr. Yogita Sharma, Dr. Rachel Fewster and Dr. Mark Bravington for discussions regarding the application of CKMR methods to mosquito populations.

## Author contributions

**Conceptualization:** John M. Marshall, Gordana Rašić

**Data curation:** John M. Marshall, Shuyi Yang, Jared B. Bennett, Igor Filipović, Gordana Rašić

**Formal analysis:** John M. Marshall, Shuyi Yang, Jared B. Bennett, Igor Filipović, Gordana Rašić

**Funding acquisition:** John M. Marshall, Gordana Rašić

**Investigation:** John M. Marshall, Shuyi Yang, Igor Filipović, Gordana Rašić

**Methodology:** John M. Marshall, Shuyi Yang, Jared B. Bennett, Igor Filipović, Gordana Rašić

**Project administration:** John M. Marshall

**Resources:** John M. Marshall, Gordana Rašić

**Software:** John M. Marshall, Shuyi Yang, Jared B. Bennett

**Supervision:** John M. Marshall

**Validation:** John M. Marshall

**Visualization:** John M. Marshall, Shuyi Yang, Gordana Rašić

**Writing - original draft preparation:** John M. Marshall

**Writing - review & editing:** John M. Marshall, Shuyi Yang, Jared B. Bennett, Igor Filipović, Gordana Rašić

## Funding statement

This work was supported by a National Institutes of Health R01 grant (1R01AI143698) awarded to J.M.M. and G.R.

## Competing interests

The authors have declared that no competing interests exist.

## Data availability

The source code for the individual-based mosquito simulation model is available at https://github.com/GilChrist19/mPlex. Documentation, including vignettes, are included for all simulation functions. The source code for inferring parameters based on the likelihood of the kinship data is available at https://github.com/MarshallLab/CKMR. Both sets of code are available under the GPL3 License and are free for other groups to modify and extend as needed.

## Notes

https://github.com/MarshallLab/CKMR

### Competing Interest Statement

The authors have declared no competing interest.

