## Supplementary material for "Spatial close-kin mark-recapture methods to estimate dispersal parameters and barrier strength for mosquitoes": S1 Text

### Spatial close-kin mark-recapture methods to estimate dispersal parameters and barrier strength for mosquitoes
### S1 Text: Supplemental model equations

John M. Marshall[1,2*], Shuyi Yang[1], Jared B. Bennett[1], Igor Filipović[3], and Gordana Rašić[3]

[1]Divisions of Biostatistics and Epidemiology, School of Public Health, University of California, Berkeley, CA, USA
[2]Innovative Genomics Institute, University of California, Berkeley, CA, USA
[3]Mosquito Genomics, QIMR Berghofer Medical Research Institute, Brisbane, Australia

May 2025

#### 1 Dispersal dynamics:

In the mosquito dispersal literature, most measurements of dispersal are over a lifetime, whereas our models are parameterized with daily mean dispersal estimates. Here, based on the theory of diffusion in two dimensions, we derive mean daily dispersal distance as a function of mean lifetime dispersal distance. We first consider dispersal in continuous space and then consider dispersal through a metapopulation.

##### 1.1 Continuous space:

In two-dimensional diffusion theory, mean displacement as a function of time, $t$, is given by:

$$x(t) = 2\sqrt{Dt}. \tag{1}$$

Here, $D$ is the diffusion constant. For diffusion in time units of days, mean daily dispersal is therefore:

$$x_d = 2\sqrt{D}. \tag{2}$$

Assuming that adult mosquitoes die according to a daily mortality rate, $\mu_A$, mosquito lifetime is exponentially distributed according to:

$$\mu_A e^{-\mu_A t}. \tag{3}$$

Expected lifetime dispersal is therefore given by the expectation:

$$x_l = \int_0^\infty \mu_A e^{-\mu_A t} \times 2\sqrt{Dt}\ dt = \sqrt{\frac{\pi D}{\mu_A}}. \tag{4}$$

Rearranging Equation 4, we have a formula for $D$ in terms of $x_l$:

$$D = \frac{x_l^2 \mu_A}{\pi}. \tag{5}$$

Finally, substituting this into Equation 2, we have the following formula for mean daily dispersal distance in terms of mean lifetime dispersal distance:

$$x_d = 2x_l\sqrt{\frac{\mu_A}{\pi}}. \tag{6}$$

### 2 Kinship probabilities:

Here, we include the kinship probabilities for parent-offspring and full-sibling pairs that, for brevity, were not included in the manuscript (§2.2).

#### 2.1 Mother-offspring:

First, we consider the mother-adult offspring kinship probability, $P_{MOA}(x_2, t_2|x_1, t_1)$, which represents the probability that, given an adult female sampled at location $x_1$ on day $t_1$, an adult sampled at location $x_2$ on day $t_2$ is her offspring. This can be expressed as the relative adult reproductive output at location $x_2$ on day $t_2$ of an adult female sampled at location $x_1$ on day $t_1$:

$$\begin{aligned} P_{MOA}(x_2, t_2|x_1, t_1) = \frac{\mathbb{E}\left[\text{Adult offspring at } (x_2, t_2) \text{ from an adult female sampled at } (x_1, t_1)\right]}{\mathbb{E}\left[\text{Adult offspring at } (x_2, t_2) \text{ from all adult females}\right]} \\ = \frac{E_{MOA}(x_2, t_2|x_1, t_1)}{E_A(x_2)}. \end{aligned} \tag{7}$$

Here, $E_{MOA}(x_2, t_2|x_1, t_1)$ represents the expected number of surviving adult offspring at location $x_2$ on day $t_2$ from an adult female sampled at location $x_1$ on day $t_1$, and $E_A(x_2)$ is given in Equation 9 of the manuscript. $E_{MOA}(x_2, t_2|x_1, t_1)$ is then given by:

$$\begin{aligned} E_{MOA}(x_2, t_2|x_1, t_1) = \sum_{y_2=t_2-T_E-T_L-T_P-(T_A-1)}^{t_2-T_E-T_L-T_P} (1-\mu_A)^{(t_1-y_2)} \\ \times \left( \begin{aligned} \mathbb{I}[(t_1 - T_A) < y_2 \leq t_1] \times \psi_{MOA}(x_2, t_2|x_1, t_1, y_2) \times \beta \times (1-\mu_E)^{T_E} \\ \times (1-\mu_L)^{T_L} \times (1-\mu_P)^{T_P} \times (1-\mu_A)^{(t_2-y_2-T_E-T_L-T_P)} \end{aligned} \right). \end{aligned} \tag{8}$$

Here, the day of egg-laying, $y_2$, is summed over days $(t_2 - T_E - T_L - T_P - (T_A - 1))$ through $(t_2 - T_E - T_L - T_P)$, for consistency with the adult offspring being present on the day of sampling (Figure **S1**). The terms within the summation are the same as for the mother-larval offspring case, with the exception that: i) daily egg production is multiplied by the proportion of eggs that survive the egg, larva, pupa and adult stages from the day they were laid up to the day of sampling, $t_2$, which reflects the additional time elapsed for adult sampling, and ii) there are now two movement types that can occur - the mother can move between egg-laying and sampling, and the adult offspring can move between emergence and sampling. These movements are captured by the composite movement term, $\psi_{MOA}(x_2, t_2|x_1, t_1, y_2)$, which represents the probability that the adult offspring is sampled at location $x_2$ on day $t_2$ given that the mother is sampled at location $x_1$ on day $t_1$ and the egg is laid on day $y_2$.

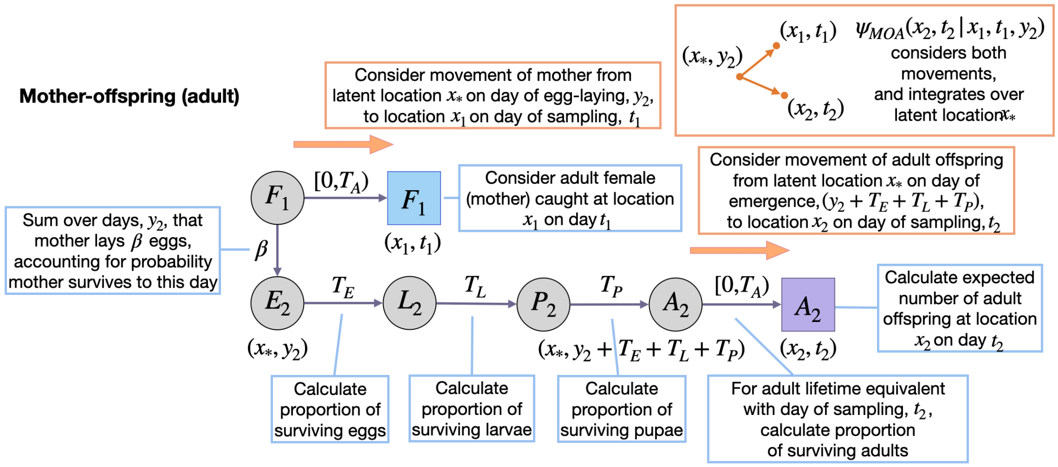


**Figure S1**: **Schematic representation of spatial mother-adult offspring kinship probability.** Parameters and state variables are as defined in Table 1 and §2.1 of the manuscript. Subscript 1 refers to the parent, and subscript 2 refers to the offspring (the perspective from which probabilities are calculated). Circles represent living individuals, squares represent sampled individuals, and colors represent their locations: blue for the sampled parent, $x_1$, purple for the sampled offspring, $x_2$, and grey for the location of egg-laying, $x_*$. Parents are sampled on day $t_1$, eggs are laid on day $y_2$, and offspring are sampled on day $t_2$. Offspring kinship probabilities are the ratio of the expected number of surviving offspring from a given adult at location $x_2$ on day $t_2$ (shown), and the expected number of surviving offspring from all adult females for this location and day (not shown). Calculating the expected number of surviving adult offspring at location $x_2$ on day $t_2$ requires considering days of egg-laying, $y_2$, consistent with maternal and adult offspring ages at sampling in the range $[0, T_A)$. There are two movements to consider: those of the mother and adult offspring (orange arrows).

Calculating $\psi_{MOA}(x_2, t_2|x_1, t_1, y_2)$ requires taking an expectation over a latent egg-laying location, $x_*$, and multiplying the probability that the offspring is laid as an egg at location $x_*$, given that the mother is sampled at location $x_1$, by the probability that the adult offspring is sampled at location $x_2$, given that it is laid as an egg at location $x_*$ (the former probability requires normalizing over all egg-laying locations), i.e.:

$$\psi_{MOA}(x_2, t_2|x_1, t_1, y_2) = \sum_{x_*} \left( \frac{\rho(x_1, t_1|x_*, y_2)}{\sum_{x_i} \rho(x_1, t_1|x_i, y_2)} \right) \times \rho(x_2, t_2|x_*, (y_2 + T_E + T_L + T_P)). \quad (9)$$

#### 2.2 Father-offspring:

Next, we consider the father-larval offspring kinship probability, $P_{FOL}(x_2, t_2|x_1, t_1)$, which represents the probability that, given an adult male sampled at location $x_1$ on day $t_1$, a larva sampled at location $x_2$ on day $t_2$ is his offspring. This can be expressed as the relative larval reproductive output at location $x_2$ on day $t_2$ of adult females that mated with an adult male sampled at location $x_1$ on day $t_1$:

$$P_{FOL}(x_2, t_2|x_1, t_1) = \frac{\mathbb{E}\left[\text{Larval offspring at } (x_2, t_2) \text{ from an adult male sampled at } (x_1, t_1)\right]}{\mathbb{E}\left[\text{Larval offspring at } (x_2, t_2) \text{ from all adult females}\right]} = \frac{E_{FOL}(x_2, t_2|x_1, t_1)}{E_L(x_2)}. \tag{10}$$

Here, $E_{FOL}(x_2, t_2|x_1, t_1)$ represents the expected number of surviving larval offspring at location $x_2$ on day $t_2$ of an adult male sampled at location $x_1$ on day $t_1$, and $E_L(x_2)$ is given as Equation 4 in the manuscript. Each adult female mates once upon emergence and, since there are equal numbers of adult females and males in the population, each adult male mates on average once in their lifetime too. The day of this mating event, $t_i$, is unknown and so, in calculating $E_{FOL}(x_2, t_2|x_1, t_1)$, we treat this as a latent variable and take an expectation over all possible values it can take:

$$E_{FOL}(x_2, t_2|x_1, t_1) = \sum_{t_i = t_1 - (T_A - 1)}^{t_1} p_A(t_1 - t_i) \times E_{FOL}(x_2, t_2|x_1, t_1, t_i). \tag{11}$$

Here, the expectation over the day of mating, $t_i$, is taken over days $(t_1 - (T_A - 1))$ through $t_1$, for consistency with the day of adult male sampling. The term $E_{FOL}(x_2, y_2|x_1, t_1, t_i)$ represents the expected number of surviving larval offspring at location $x_2$ on day $t_2$ of an adult male sampled at location $x_1$ on day $t_1$, conditional upon the day of mating being $t_i$, and is given by:

$$E_{FOL}(x_2, t_2|x_1, t_1, t_i) = \sum_{y_2 = t_i}^{t_i + (T_A - 1)} (1 - \mu_A)^{(y_2 - t_i)} \times \begin{pmatrix} \mathbb{I}[(y_2 + T_E) \leq t_2 < (y_2 + T_E + T_L)] \\ \times \psi_{FOL}(x_2, t_2|x_1, t_1, t_i, y_2) \times \beta \\ \times (1 - \mu_E)^{T_E} \times (1 - \mu_L)^{(t_2 - y_2 - T_E)} \end{pmatrix}. \tag{12}$$

Here, the day of egg-laying, $y_2$, is summed over days $t_i$ through $(t_i + (T_A - 1))$, for consistency with the mother's potential lifespan. The terms within the summation are then the same as for the mother-larval offspring case, with the exception that: i) the indicator function limits consideration to cases where the day of larval sampling lies within the larval offspring's possible lifetime - i.e. between days $(y_2 + T_E)$ and $(y_2 + T_E + (T_L - 1))$, and ii) the movement of interest is now of the mother between egg-laying events. This is captured by the movement term, $\psi_{FOL}(x_2, t_2|x_1, t_1, t_i, y_2)$, which represents the movement probability corresponding to the larval offspring being sampled at location $x_2$ on day $t_2$ given that the father is sampled at location $x_1$ on day $t_1$, the day of mating is $t_i$, and the day of egg-laying is $y_2$.

Calculating $\psi_{FOL}(x_2, t_2|x_1, t_1, t_i, y_2)$ requires considering the father's movement between mating and sampling, in addition to the mother's movement between mating and egg-laying, i.e.:

$$\psi_{FOL}(x_2, t_2|x_1, t_1, t_i, y_2) = \sum_{x_i} \left( \frac{\rho(x_1, t_1|x_i, t_i)}{\sum_{x_j} \rho(x_1, t_1|x_j, t_i)} \right) \times \rho(x_2, y_2|x_i, t_i). \tag{13}$$

Here, we take an expectation over a latent mating location, $x_i$, and multiply the probability that mating happens at location $x_i$ given that the father is sampled at location $x_1$ by the probability that the larval offspring is sampled at location $x_2$ (the former probability requires normalizing over all possible mating locations).

Next, we consider the father-adult offspring kinship probability, $P_{FOA}(x_2, t_2|x_1, t_1)$, which represents the probability that, given an adult male sampled at location $x_1$ on day $t_1$, an adult sampled at location $x_2$ on day $t_2$ is his offspring. This can be expressed as the relative adult reproductive output at location $x_2$ on day $t_2$ of adult females that mated with an adult male sampled at location $x_1$ on day $t_1$:

$$\begin{aligned}P_{FOA}(x_2, t_2|x_1, t_1) = \frac{\mathbb{E}\left[\text{Adult offspring at } (x_2, t_2) \text{ from an adult male sampled at } (x_1, t_1)\right]}{\mathbb{E}\left[\text{Adult offspring at } (x_2, t_2) \text{ from all adult females}\right]} \\ = \frac{E_{FOA}(x_2, t_2|x_1, t_1)}{E_A(x_2)}.\end{aligned} \tag{14}$$

Here, $E_{FOA}(x_2, t_2|x_1, t_1)$ represents the expected number of surviving adult offspring at location $x_2$ on day $t_2$ of an adult male sampled at location $x_1$ on day $t_1$, and $E_A(x_2)$ is given in Equation 9 of the manuscript. Each adult male mates on average once; but the day of this mating event, $t_i$, is unknown and so, in calculating $E_{FOA}(x_2, t_2|x_1, t_1)$, we treat this as a latent variable and take an expectation over all possible values it can take:

$$E_{FOA}(x_2, t_2|x_1, t_1) = \sum_{t_i=t_1-(T_A-1)}^{t_1} p_A(t_1 - t_i) \times E_{FOA}(x_2, t_2|x_1, t_1, t_i). \tag{15}$$

Here, the expectation over the day of mating, $t_i$, is taken over days $(t_1 - (T_A - 1))$ through $t_1$, for consistency with the day of adult male sampling. The term $E_{FOA}(x_2, t_2|x_1, t_1, t_i)$ represents the expected number of adult offspring at location $x_2$ on day $t_2$ of an adult male sampled at location $x_1$ on day $t_1$, conditional upon the day of mating being $t_i$, and $p_A(t_1 - t_i)$ represents the probability that the mating event occurs on day $(t_1 - t_i)$. $E_{FOA}(x_2, t_2|x_1, t_1, t_i)$ is then given by:

$$\begin{aligned}E_{FOA}(x_2, t_2|x_1, t_1, t_i) = \sum_{y_2=t_i}^{t_i+(T_A-1)} (1-\mu_A)^{(y_2-t_i)} \\ \times \begin{pmatrix}\mathbb{I}[(y_2 + T_E + T_L + T_P) \leq t_2 < (y_2 + T_E + T_L + T_P + T_A)] \\ \times \psi_{FOA}(x_2, t_2|x_1, y_1, t_i) \times \beta \times (1-\mu_E)^{T_E} \times (1-\mu_L)^{T_L} \\ \times (1-\mu_P)^{T_P} \times (1-\mu_A)^{(t_2-y_2-T_E-T_L-T_P)}\end{pmatrix}.\end{aligned} \tag{16}$$

Here, the day of egg-laying, $y_2$, is summed over days $t_i$ through $(t_i + (T_A - 1))$, for consistency with the mother's potential lifespan. The first term in the summation represents the probability that the mother is alive on the day of egg-laying, and the second term (in larger brackets) represents the expected surviving adult output of this adult female on day $t_2$. This latter term is the same as for the mother-adult offspring case, with the exception that: i) the indicator function limits consideration to cases where the day of adult sampling, $t_2$, lies within the adult offspring's possible lifetime - i.e. between days $(y_2 + T_E + T_L + T_P)$ and $(y_2 + T_E + T_L + T_P + (T_A - 1))$, and ii) there are now three movement events that can occur - the father can move between mating and sampling, the mother can move between mating and egg-laying, and the adult offspring can move been emergence and sampling. These movements are captured by the composite movement term, $\psi_{FOA}(x_2, t_2|x_1, y_1, t_i)$, which represents the movement probability corresponding to the adult offspring being sampled at location $x_2$ on day $t_2$ given that the father is sampled at location $x_1$ on day $t_1$, and the day of mating is $t_i$.

Calculating $\psi_{FOA}(x_2, t_2|x_1, t_1, t_i)$ requires considering the father's movement between mating and sampling, in addition to the mother's movement between mating and egg-laying, and the adult offspring's movement between emergence and sampling, i.e.:

$$\psi_{FOA}(x_2, t_2|x_1, t_1, t_i) = \sum_{x_i} \left( \frac{\rho(x_1, t_1|x_i, t_i)}{\sum_{x_j} \rho(x_1, t_1|x_j, t_i)} \right) \times \rho(x_2, t_2|x_i, (t_i + T_E + T_L + T_P)). \tag{17}$$

Here, we take an expectation over a latent mating location, $x_i$, and multiply the probability that mating happens at location $x_i$ given that the father is sampled at location $x_1$ by the probability that the adult offspring is sampled at location $x_2$ (the former probability requires normalizing over all mating locations).

##### 2.3 Full-siblings:

Next, we consider the full-sibling kinship probability for larva-larva pairs, $P_{FSLL}(x_2, t_2|x_1, t_1)$, which represents the probability that, given a larva sampled at location $x_1$ on day $t_1$, a larva sampled at location $x_2$ on day $t_2$ is their full-sibling. This can be expressed as the relative larval reproductive output at location $x_2$ on day $t_2$ of the mother of a larva sampled at location $x_1$ on day $t_1$:

$$\begin{aligned} P_{FSLL}(x_2, t_2|x_1, t_1) &= \frac{\mathbb{E}\left[\text{Larvae at } (x_2, t_2) \text{ that are full-siblings of a larva sampled at } (x_1, t_1)\right]}{\mathbb{E}\left[\text{Larval offspring at } (x_2, t_2) \text{ from all adult females}\right]} \\ &= \frac{E_{FSLL}(x_2, t_2|x_1, t_1)}{E_L(x_2)}. \end{aligned} \tag{18}$$

Here, $E_{FSLL}(x_2, t_2|x_1, t_1)$ represents the expected number of surviving larvae at location $x_2$ on day $t_2$ that are full-siblings of a larva sampled at location $x_1$ on day $t_1$, and $E_L$ is given in Equation 4 of the manuscript. For convenience, let us refer to the larva sampled at location $x_1$ on day $t_1$ as individual 1. To calculate $E_{FSLL}(x_2, t_2|x_1, t_1)$, we treat the day that egg 1 is laid, $y_1$, as a latent variable and take an expectation over it:

$$E_{FSLL}(x_2, t_2|x_1, t_1) = \sum_{y_1 = t_1 - T_E - (T_L - 1)}^{t_1 - T_E} p_L(t_1 - y_1 - T_E) \times E_{FSLL}(x_2, t_2|x_1, t_1, y_1). \tag{19}$$

Here, the expectation over the day that egg 1 is laid, $y_1$, is taken over days $(t_1 - T_E - (T_L - 1))$ through $(t_1 - T_E)$, for consistency with the day that larva 1 is sampled (Figure **S2**). The term $E_{FSLL}(x_2, t_2|x_1, t_1, y_1)$ represents the expected number of surviving larvae at location $x_2$ on day $t_2$ that are full-siblings of larva 1, conditional upon egg 1 being laid on day $y_1$. Additionally, $p_L(t_1 - y_1 - T_E)$ represents the probability that egg 1 is laid on day $(t_1 - y_1 - T_E)$. In general, $p_L(t)$ represents the probability that a given larva in the population has age $t$ which, following from the daily larval survival probability, $(1 - \mu_L)$, is given by:

$$p_L(t) = (1 - \mu_L)^t \Big/ \sum_{t_j=0}^{T_L - 1} (1 - \mu_L)^{t_j}. \tag{20}$$

$E_{FSLL}(x_2, t_2|x_1, t_1, y_1)$ is then given by:

$$\begin{aligned} E_{FSLL}(x_2, t_2|x_1, t_1, y_1) = \frac{1}{2} &\sum_{y_2 = y_1 - (T_A - 1)}^{y_1 + (T_A - 1)} (1 - \mu_A)^{|y_2 - y_1|} \\ \times &\begin{pmatrix} \mathbb{I}[(t_2 - T_E - T_L) < y_2 \leq (t_2 - T_E)] \times \psi_{FSLL}(x_2, t_2|x_1, t_1, y_1, y_2) \\ \times \beta \times (1 - \mu_E)^{T_E} \times (1 - \mu_L)^{(t_2 - y_2 - T_E)} \end{pmatrix}. \end{aligned} \tag{21}$$

Here, the day of sibling egg-laying, $y_2$, is summed over days $y_1 - (T_A - 1)$ through $y_1 + (T_A - 1)$, for consistency with the mother's potential lifespan (Figure **S2**), and considering that either of the larval offspring may have been laid first. Since we are summing two equally-weighted scenarios regarding offspring order,

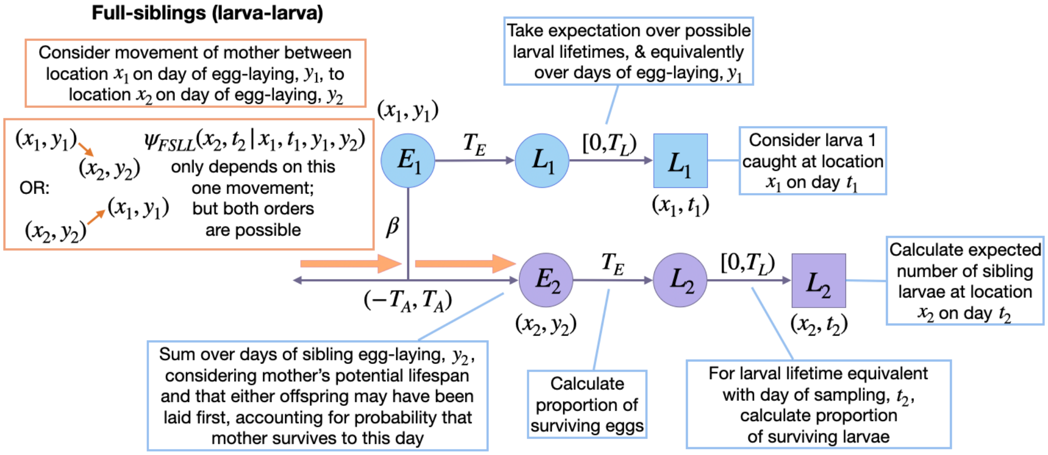


**Figure S2**: **Schematic representation of spatial larva-larva full-sibling kinship probabilities.** Parameters and state variables are as defined in Table 1 and §2.2 of the manuscript. Subscript 1 refers to the reference sibling, and subscript 2 refers to the sibling from whose perspective the probabilities are calculated. Circles represent living individuals, squares represent sampled individuals, and colors represent their locations: blue for sibling 1, $x_1$, and purple for sibling 2, $x_2$. Sibling 1 is sampled on day $t_1$ and laid on day $y_1$. Sibling 2 is sampled on day $t_2$ and laid on day $y_2$. Sibling kinship probabilities are the ratio of the expected number of surviving siblings of a given individual at location $x_2$ on day $t_2$ (shown), and the expected number of surviving offspring from all adult females for this location and day (not shown). Calculating the expected number of surviving full-siblings at location $x_2$ on day $t_2$ requires considering days of egg-laying, $y_1$ and $y_2$, consistent with adult ages at sampling in the range $[0, T_A)$. The only movement to consider is that of the mother between egg-laying events. Movement probabilities consider both orders of egg-laying (orange arrows).

we include a multiplier of 1/2 in the expectation. The first term within the summation then represents the probability that the mother is alive on the day of sibling egg-laying, with the absolute value, $|y_2 - y_1|$, accounting for both offspring orders. The second term (in larger brackets) represents the expected larval output of the mother of larva 1 at location $x_2$ on day $t_2$. This latter term is the same as for the mother-larval offspring case, with the exception that: i) the indicator function limits consideration to cases where the day of sibling egg-laying, $y_2$, is between days $(t_2 - T_E - (T_L - 1))$ and $(t_2 - T_E)$, for consistency with a larval sibling being sampled on day $t_2$, and ii) the movement of interest is now of the mother between egg-laying events. This is captured by the movement term, $\psi_{FSLL}(x_2, t_2 | x_1, t_1, y_1, y_2)$, which represents the probability that larval offspring 2 is sampled at location $x_2$ on day $t_2$ given that larval offspring 1 is sampled at location $x_1$ on day $t_1$.

Calculating $\psi_{FSLL}(x_2, t_2 | x_1, t_1, y_1, y_2)$ requires considering the mother's movement between egg-laying events for both offspring orders, i.e.:

$$\psi_{FSLL}(x_2, t_2 | x_1, t_1, y_1, y_2) = \begin{cases} 1_{x_1 = x_2} & , y_2 = y_1 \\ \rho(x_2, y_2 | x_1, y_1) & , y_2 > y_1 \\ \frac{\rho(x_1, y_1 | x_2, y_2)}{\sum_{x_i} \rho(x_1, y_1 | x_i, y_2)} & , y_2 < y_1 \end{cases} . \tag{22}$$

In the case where both offspring are laid on the same day, no movement occurs; in the case where offspring 1 is laid first, the movement probability defined in Equation 7 directly applies to the egg-laying events; and in the case where offspring 2 is laid first, given that the mother is at location $x_1$ at time $y_1$, the probability that her previous location at time $y_2$ is $x_2$ is calculated by normalizing over all possible egg-laying locations.

Next, we consider the larva-adult full-sibling kinship probability, $P_{FSLA}(x_2, t_2|x_1, t_1)$, which represents the probability that, given a larva sampled at location $x_1$ on day $t_1$, an adult sampled at location $x_2$ on day $t_2$ is their full-sibling:

$$\begin{aligned} P_{FSLA}(x_2, t_2|x_1, t_1) &= \frac{\mathbb{E}\left[\text{Adults at } (x_2, t_2) \text{ that are full-siblings of a larva sampled at } (x_1, t_1)\right]}{\mathbb{E}\left[\text{Adult offspring at } (x_2, t_2) \text{ from all adult females}\right]} \\ &= \frac{E_{FSLA}(x_2, t_2|x_1, t_1)}{E_A(x_2)}. \end{aligned} \tag{23}$$

Here, $E_{FSLA}(x_2, t_2|x_1, t_1)$ represents the expected number of surviving adults at location $x_2$ on day $t_2$ that are full-siblings of a larva sampled at location $x_1$ on day $t_1$, and $E_A(x_2)$ is given in Equation 9 of the manuscript. For convenience, let us refer to the larva sampled at location $x_1$ on day $t_1$ as individual 1. To calculate $E_{FSLA}(x_2, t_2|x_1, t_1)$, we treat the day that egg 1 is laid, $y_1$, as a latent variable and take an expectation over it:

$$E_{FSLA}(x_2, t_2|x_1, t_1) = \sum_{y_1=t_1-T_E-(T_L-1)}^{t_1-T_E} p_L(t_1 - y_1 - T_E) \times E_{FSLA}(x_2, t_2|x_1, t_1, y_1). \tag{24}$$

This is the same equation as for the larva-larva case with the only exception being the term $E_{FSLA}(x_2, t_2|x_1, t_1, y_1)$, which represents the expected number of surviving adults at location $x_2$ on day $t_2$ that are full-siblings of larva 1, conditional upon egg 1 being laid on day $y_1$. That term is given by:

$$\begin{aligned} E_{FSLA}(x_2, t_2|x_1, t_1, y_1) = \frac{1}{2} \sum_{y_2=y_1-(T_A-1)}^{y_1+(T_A-1)} (1-\mu_A)^{|y_2-y_1|} \\ \times \begin{pmatrix} \mathbb{I}[(t_2 - T_E - T_L - T_P - T_A) < y_2 \leq (t_2 - T_E - T_L - T_P)] \\ \times \psi_{FSLA}(x_2, t_2|x_1, t_1, y_1, y_2) \times \beta \times (1-\mu_E)^{T_E} \times (1-\mu_L)^{T_L} \\ \times (1-\mu_P)^{T_P} \times (1-\mu_A)^{(t_2-y_2-T_E-T_L-T_P)} \end{pmatrix}. \end{aligned} \tag{25}$$

This is the same equation as for the adult-adult case with the only exception being the term $\psi_{FSLA}(x_2, t_2|x_1, t_1, y_1, y_2)$, which represents the movement probability corresponding to adult offspring 2 being sampled at location $x_2$ on day $t_2$ given that larval offspring 1 is sampled at location $x_1$ on day $t_1$, egg 1 is laid on day $y_1$, and egg 2 is laid on day $y_2$. Calculating $\psi_{FSLA}(x_2, t_2|x_1, t_1, y_1, y_2)$ requires considering two movements - that of the mother between egg-laying events, and the movement of adult offspring 2 between emergence and sampling - for both offspring orders. Beginning with the case where offspring 1 is laid first or on the same day as offspring 2 (i.e., $y_2 \geq y_1$), we have:

$$\psi_{FSLA}(x_2, t_2|x_1, t_1, y_1, y_2) = \rho(x_2, t_2|x_1, (y_1 + T_E + T_L + T_P)). \tag{26}$$

Here, movements are only modeled forwards in time. For the adult sibling, movement occurs after development through the egg, larva and pupa life stages (i.e., after $(T_E + T_L + T_P)$ days), and the mother's movement between egg-laying events is incorporated by adding $(y_2 - y_1)$ days of movement to that of the adult sibling. Second, for the case where offspring 2 is laid first (i.e., $y_2 < y_1$), we have:

$$\psi_{FSLA}(x_2, t_2|x_1, t_1, y_1, y_2) = \sum_{x_*} \left( \frac{\rho(x_1, y_1|x_*, y_2)}{\sum_{x_i} \rho(x_1, y_1|x_i, y_2)} \right) \times \rho(x_2, t_2|x_*, (y_2 + T_E + T_L + T_P)). \tag{27}$$

Here, we take an expectation over a latent egg-laying location for the firstborn sibling, $x_*$, and multiply the probability that the firstborn sibling is laid at location $x_*$ given that adult sibling 1 is sampled at location $x_1$ by the probability that adult sibling 2 is sampled at location $x_2$ (the former probability requires normalizing over all egg-laying locations). For the adult sibling, movement is modeled forwards in time and begins after development through the egg, larva and pupa life stages (i.e., after $(T_E + T_L + T_P)$ days).

Next, we consider the adult-larva full-sibling kinship probability, $P_{FSAL}(x_2, t_2|x_1, t_1)$, which represents the probability that, given an adult sampled at location $x_1$ on day $t_1$, a larva sampled at location $x_2$ on day $t_2$ is their full-sibling:

$$P_{FSAL}(x_2, t_2|x_1, t_1) = \frac{\mathbb{E}\left[\text{Larvae at } (x_2, t_2) \text{ that are full-siblings of an adult sampled at } (x_1, t_1)\right]}{\mathbb{E}\left[\text{Larval offspring at } (x_2, t_2) \text{ from all adult females}\right]} = \frac{E_{FSAL}(x_2, t_2|x_1, t_1)}{E_L(x_2)}. \tag{28}$$

Here, $E_{FSAL}(x_2, t_2|x_1, t_1)$ represents the expected number of surviving larvae at location $x_2$ on day $t_2$ that are full-siblings of an adult sampled at location $x_1$ on day $t_1$, and $E_L(x_2)$ is given by Equation 4 in the manuscript. For convenience, let us refer to the adult sampled at location $x_1$ on day $t_1$ as individual 1. To calculate $E_{FSAL}(x_2, t_2|x_1, t_1)$, we treat the day that egg 1 is laid, $y_1$, as a latent variable and take an expectation over it:

$$E_{FSAL}(x_2, t_2|x_1, t_1) = \sum_{y_1 = t_1 - T_E - T_L - T_P - (T_A - 1)}^{t_1 - T_E - T_L - T_P} p_A(t_1 - y_1 - T_E - T_L - T_P) \times E_{FSAL}(x_2, t_2|x_1, t_1, y_1). \tag{29}$$

This is the same equation as for the adult-adult case with the only exception being the term $E_{FSAL}(x_2, t_2|x_1, t_1, y_1)$, which represents the expected number of surviving larvae at location $x_2$ on day $t_2$ that are full-siblings of adult 1, conditional upon egg 1 being laid on day $y_1$. That term is given by:

$$E_{FSAL}(x_2, t_2|x_1, t_1, y_1) = \frac{1}{2} \sum_{y_2 = y_1 - (T_A - 1)}^{y_1 + (T_A - 1)} (1 - \mu_A)^{|y_2 - y_1|} \times \begin{pmatrix} \mathbb{I}[(t_2 - T_E - T_L) < y_2 \leq (t_2 - T_E)] \times \psi_{FSAL}(x_2, t_2|x_1, t_1, y_1, y_2) \\ \times \beta \times (1 - \mu_E)^{T_E} \times (1 - \mu_L)^{(t_2 - y_2 - T_E)} \end{pmatrix}. \tag{30}$$

This is the same equation as for the larva-larva case with the only exception being the term $\psi_{FSAL}(x_2, t_2|x_1, t_1, y_1, y_2)$, which represents the movement probability corresponding to larval offspring 2 being sampled at location $x_2$ on day $t_2$ given that adult offspring 1 is sampled at location $x_1$ on day $t_1$, egg 1 is laid on day $y_1$, and egg 2 is laid on day $y_2$. Calculating $\psi_{FSAL}(x_2, t_2|x_1, t_1, y_1, y_2)$ requires considering two movements - that of the mother between egg-laying events, and the movement of adult offspring 1 between emergence and sampling - for both offspring orders. Beginning with the case where offspring 2 is laid first or on the same day as offspring 1 (i.e., $y_2 \leq y_1$), we have:

$$\psi_{FSAL}(x_2, t_2|x_1, t_1, y_1, y_2) = \frac{\rho(x_1, t_1|x_2, (y_2 + T_E + T_L + T_P))}{\sum_{x_i} \rho(x_1, t_1|x_i, (y_2 + T_E + T_L + T_P))}. \tag{31}$$

Here, movements are only modeled backwards in time. For the adult sibling, movement occurs after development through the egg, larva and pupa life stages (i.e., after $(T_E + T_L + T_P)$ days), and the mother's movement between egg-laying events is incorporated by adding $(y_2 - y_1)$ days of movement to that of the adult sibling. Second, for the case where offspring 1 is laid first (i.e., $y_2 > y_1$), we have:

$$\psi_{FSAL}(x_2, t_2|x_1, t_1, y_1, y_2) = \sum_{x_*} \left( \frac{\rho(x_1, t_1|x_*, (y_1 + T_E + T_L + T_P))}{\sum_{x_i} \rho(x_1, t_1|x_i, (y_1 + T_E + T_L + T_P))} \right) \times \rho(x_2, y_2|x_*, y_1). \tag{32}$$

Here, we take an expectation over a latent egg-laying location for the firstborn sibling, $x_*$, and multiply the probability that the firstborn sibling is laid at location $x_*$ given that adult sibling 1 is sampled at location $x_1$ by the probability that larval sibling 2 is sampled at location $x_2$ (the former probability requires normalizing over all egg-laying locations). For the mother, movement is modeled forwards in time and occurs between egg-laying events (i.e., after between days $y_1$ and $y_2$).
